# Systematic mapping of drug metabolism by the human gut microbiome

**DOI:** 10.1101/538215

**Authors:** Pranatchareeya Chankhamjon, Bahar Javdan, Jaime Lopez, Raphaella Hull, Seema Chatterjee, Mohamed S. Donia

## Abstract

The human gut microbiome harbors hundreds of bacterial species with diverse biochemical capabilities, making it one of nature’s highest density, highest diversity bioreactors. Several drugs have been previously shown to be directly metabolized by the gut microbiome, but the extent of this phenomenon has not been systematically explored. Here, we develop a systematic screen for mapping the ability of the complex human gut microbiome to biochemically transform small molecules (MDM-Screen), and apply it to a library of 575 clinically used oral drugs. We show that 13% of the analyzed drugs, spanning 28 pharmacological classes, are metabolized by a single microbiome sample. In a proof-of-principle example, we show that microbiome-derived metabolism occurs *in vivo*, identify the genes responsible for it, and provide a possible link between its consequences and clinically observed features of drug bioavailability and toxicity. Our findings reveal a previously underappreciated role for the gut microbiome in drug metabolism, and provide a comprehensive framework for characterizing this important class of drug-microbiome interactions.

## INTRODUCTION

The oral route is the most common route for drug administration. Upon exiting the stomach, drugs can be absorbed in the small and/or large intestine to reach the systemic circulation and eventually the liver, or can be transported there directly via the portal vein. Once at the liver, drugs may be metabolized and secreted back (along with their metabolites) to the intestines through bile, via the enterohepatic circulation^1,2^. Even parenterally administered drugs and their resulting metabolites can reach the intestines through biliary secretion. Therefore, whether prior to or after absorption, most administered drugs will spend a considerable amount of time in the small and large intestines, where trillions of bacterial cells reside and form our human gut microbiome. Despite this fact, and the significant inter-individual variability in both the composition and function of the gut microbiome^3^, we know much less about how our microbiome interacts with drugs than about how our liver interacts with them.

Broadly speaking, there are two main types of interactions that can occur between drugs and the microbiome, which may result in significant effects on drug metabolism, bioavailability, efficacy, and toxicity: direct and indirect interactions. Examples of indirect interactions include the competition between microbiome-derived metabolites and administered drugs for the same host metabolizing enzymes^4^, microbiome effects on the immune system in anticancer immunotherapy^5-7^, microbiome reactivation of secreted inactive metabolites of the drug^8^, and microbiome overall effects on the levels of metabolizing enzymes in the liver and intestine^9^. Direct interactions between administered drugs and the microbiome include the partial or complete biochemical transformation of a drug into more or less active metabolites by microbiome-derived enzymes (termed herein: Microbiome-Derived Metabolism, or MDM).

The human gut microbiome harbors hundreds of bacterial species, encoding an estimated 100 times more genes than the human genome^10^. This enormous diversity and richness of genes represent a repertoire of yet-uncharacterized biochemical activities capable of metabolizing ingested chemicals, including both dietary and therapeutic ones^11^. Although MDM has been observed for more than 50 years, and in dozens of examples, this process is still mostly overlooked in the drug development pipeline^2,12-14^. Moreover, studies investigating this process have focused mainly on one bacterium or one drug at a time, and no efforts have been spent to systematically assess the ability of the human gut microbiome to metabolize oral drugs or to develop tools for incorporating this type of analysis into the drug development pipeline. This is owed mainly to the enormous complexity of the microbiome, and to the overwhelming technical challenge of testing hundreds of drugs against thousands of cultured isolates under multiple conditions. Unlike liver-derived metabolism, the lack of a systematic, global, and standardized map of MDM has hindered our ability to reliably predict and eventually interfere with undesired microbiome effects on drug pharmacokinetics and/or pharmacodynamics.

To address this gap in knowledge, we here develop a systematic screen for mapping MDM (MDM-Screen, **Fig. 1**). Our screen relies on three main arms: i) an optimized batch culturing system for sustaining the growth of complex, personalized, human microbiome-derived microbial communities; ii) a high-throughput analytical chemistry platform for screening hundreds of clinically used small molecule drugs; and iii) a defined mouse colonization assay for assessing the effect of the microbiome on the pharmacokinetics of selected drugs. Using MDM-Screen with 575 clinically used, orally administered, small molecule drugs, we discovered that 13% of them can be subject to MDM. As a proof-of-principle example, we selected one of these transformations – MDM deglycosylation of fluoropyrimidines – for further functional investigations. We identify microbiome-derived species and enzymes responsible for this transformation, show that it occurs *in vivo* in a microbiome-dependent manner, and provide evidence that its consequences may explain outcomes already observed in the clinic. Our screen described here, and the findings obtained from it represent the first systematic map of microbiome-derived metabolism of clinically used drugs, and provide a framework for incorporating an “MDM” module in future drug development pipelines.

**Figure 1.**
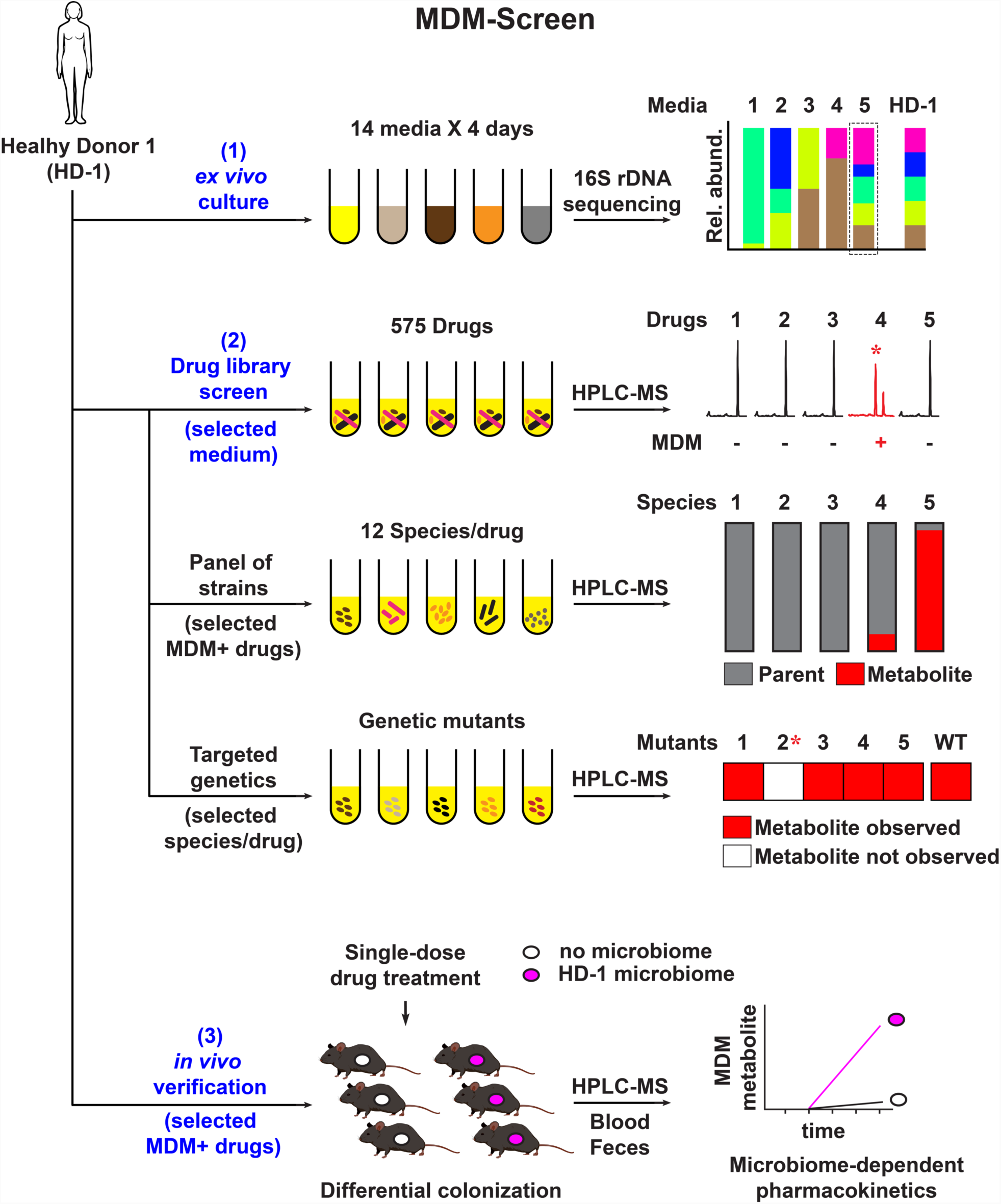
General approach of MDM-Screen. MDM-Screen is comprised of three arms. (1) An optimized *ex vivo* culturing model of the gut microbiome in batch format, where a fecal sample from a healthy donor (HD-1) is cultured in 14 different media for 4 days, and the best culturing condition is determined by high-throughput 16S rDNA amplicon sequencing. (2) A biochemical screen, where the ability of the cultured HD-1 microbiome to metabolize 575 drugs is determined using HPLC-MS. By screening a diverse set of gut isolates, the same platform is used to identify members of the microbiome that may be responsible for specific modifications. Finally, specific genes and enzymes responsible for the modifications are identified by targeted mutagenesis in selected species. (3) For selected MDM cases, a microbiome-dependent pharmacokinetic experiment is performed in mice to assess whether the same drug modification can be observed *in vivo*.

## RESULTS

### An optimized *ex vivo* culturing model for the human gut microbiome

A major challenge in studying the capacity of the human gut microbiome to metabolize orally administered drugs is the enormous diversity of the bacterial species involved: a typical gut microbiome sample harbors hundreds of species and thousands of strains, many of which are found only in a subset of healthy individuals^3,15^. It is therefore impractical to systematically screen thousands of isolated strains against hundreds of drugs, forcing previous studies to rely mainly on a selected set of representative species. Moreover, gene expression profiles and the significance of a given biochemical transformation may vary dramatically between a monocultured strain and one that is grown in a mixed community. To address these challenges, we sought to develop the first arm of MDM-Screen: an optimized *ex vivo* culturing system that a) supports the growth of a large proportion of the species from a given microbiome sample in a similar taxonomical composition, and b) is amenable to high-throughput biochemical screens.

Acknowledging the fact that a significant fraction of the community will inevitably evade cultivation efforts, we undertook a systematic approach to identify the medium and culturing period that can support the growth of the maximal number of species in a batch culture of a mixed community. Freshly collected human feces from a healthy donor (referred to as HD-1) were transferred to an anaerobic chamber, suspended in PBS with 0.1% cysteine, and stored in aliquots of dozens of glycerol stocks. We then started cultures (anaerobic, 37 °C) from glycerol-stocked HD-1 in 14 different media, and collected samples daily for 4 days. Finally, we extracted DNA from all cultures, amplified the V4 region of the bacterial 16S rRNA gene, and deeply sequenced the amplicons using Illumina (100,000 sequences per sample, on average). From the sequencing results, amplicon sequence variants (ASVs) were inferred using DADA2 plugin within QIIME2, and the final taxonomical composition at different levels was determined for each sample using a naive Bayes classifier trained on the Greengenes database^16-19^. We then quantified the differences between the various media and the original fecal sample at both the family level (using the Jensen-Shannon divergence (D_JS_), a metric that measures the similarity of two distributions), as well as at the single ASV level (to infer the recovery rate of species from the original sample).

Two main findings emerged from this analysis. First, as expected, we observed a great level of variation in both the taxonomical composition and diversity between the different media and culturing periods. Some media led to highly diverse communities that captured portions of the original fecal diversity, while others became dominated almost exclusively by a single family. Second, among the 14 media commonly used in cultivation efforts from the human microbiome^20^, we identified one medium, modified Gifu Anaerobic Medium (mGAM), that supported the growth of a bacterial community most similar in composition and diversity to the one observed in HD-1 (**Fig. 2a, Supplementary Fig. 1**). At the family level, mGAM cultures largely match the composition of HD-1, differing primarily in a commonly observed expansion of the facultative anaerobes, Enterobacteriaceae, at the expense of the obligate anaerobes, Ruminococcaceae. This is likely a result of the inevitable exposure to oxygen during sample handling until delivery to the anaerobic chamber (∼ 30 min)^21^. Among all tested media, mGAM cultures showed the lowest D_JS_ divergence from HD-1, becoming increasingly similar to the original sample as growth proceeds (see **Supplementary Fig. 1** for the entire four-day time course).

**Figure 2.**
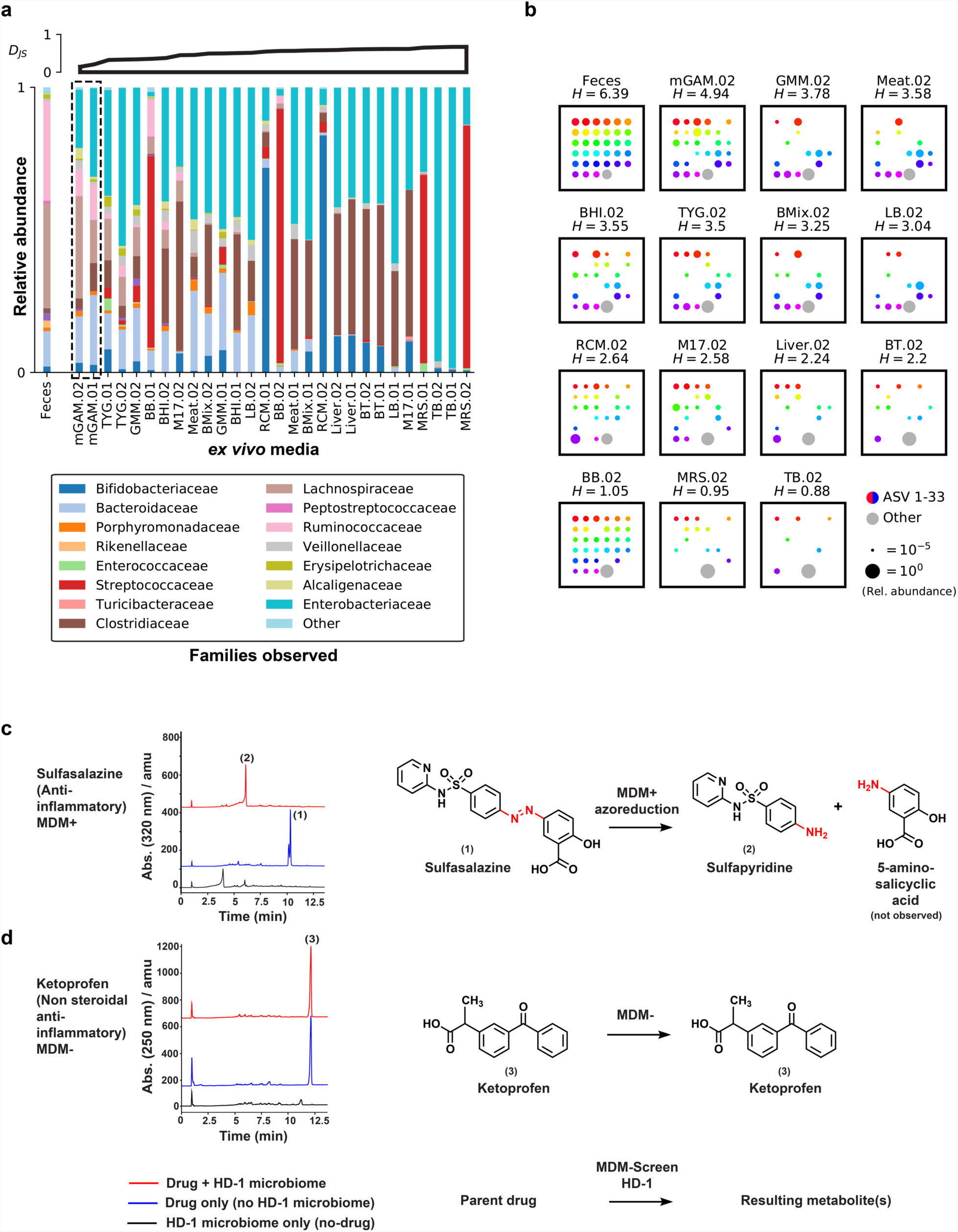
Development of MDM-Screen. **a)** Family level bacterial composition of the original HD-1 fecal sample (far left), as well as that of HD-1 *ex vivo* cultures grown anaerobically in 14 different media over two days (.01 and.02). Full names of the media used are listed in the **Methods**. A four-day time course of HD-1 in the same media is shown in **Supplementary Fig. 1**. 16S rRNA gene sequences that could not be classified at the family level, and families with less than 1% relative abundance in all samples are grouped into “Other”. Cultures are ordered according to their Jensen-Shannon D_JS_ divergence from the original HD-1 sample (upper axes, computed at the family level), where lower values indicate higher similarity to HD-1. Note that cultures grown in mGAM (mGAM.02 and mGAM.01) are the most similar to HD-1. **b)** Amplicon Sequence Variant (ASV) level bacterial composition of the original HD-1 fecal sample, and that of day two *ex vivo* cultures of HD-1 grown in 14 different media, where each square represents one sample. Rainbow colored dots represent the relative abundance of individual ASVs that are above 1% in HD-1, while grey dots represent the combined relative abundance of all ASVs below 1% in HD-1. A larger dot indicates a higher relative abundance, as indicated by a size scale at the bottom right corner. Samples are ordered by their Shannon diversity (*H*) at the ASV level, computed in bits and shown above each square. Note that mGAM.02 culture has the highest Shannon diversity, and the closest to HD-1. **c)** HPLC-MS analysis of sulfasalazine (1) incubated with HD-1 mGAM-02 culture (red) or with mGAM.02 broth (blue). A similar analysis is also done for HD-1 mGAM.02 culture with no drug added (black). An HPLC chromatogram at an absorbance of 320 nm is shown for all three samples, indicating the conversion of sulfasalazine (1) to sulfapyridine (2) in the presence of the HD-1 microbiome. This is a typical case of an MDM+ drug. **d)** A similar HPLC-MS analysis for ketoprofen (3). An HPLC chromatogram at an absorbance of 250 nm is shown, indicating no modification to the parent drug in the presence of the HD-1 microbiome. This is a typical case of an MDM-drug.

Even at the single ASV level, mGAM cultures capture much of the diversity in HD-1 (mGAM cultures have the highest Shannon diversity across all media, and the closest one to HD-1) (**Fig. 2b and Supplementary Fig. 2**). In the original fecal sample, there are 33 ASVs present above a relative abundance of 1%, 26 (79%) of which are present in mGAM day two culture. Overall, total shared ASVs between the original fecal sample and mGAM day two account for 70% of the HD-1 composition, indicating that the mGAM culture recapitulates the bulk of the original community. Taken together, and consistent with previous reports showing that mGAM can support the growth of a wide variety of gut microorganisms in monoculture^20,22^, our results establish mGAM day two cultures as a viable *ex vivo* batch culturing model for the human gut microbiome, where a significant portion of the taxonomical diversity from the original fecal sample can be captured and maintained in a similar composition.

### A high-throughput drug screen for MDM

With an optimized *ex vivo* culturing system in hand, we developed the second arm of MDM-Screen: a combined biochemical / analytical chemistry approach for the systematic mapping of MDM. Our approach needed to fulfill the following criteria: a) is reproducible, and its reproducibility can be quickly assessed, b) is scalable to hundreds of drugs, c) is sensitive, even with a small amount of drug, and d) is feasible in a reasonable time frame and in an academic laboratory setting. After several iterations, we successfully devised a strategy that meets all four desired criteria (**Fig.1 and Fig.2b, 2c**). In this strategy, three samples are prepared per drug of interest: 1) a 3-ml, 24-hour mGAM *ex vivo* culture of the starting human feces, incubated with the drug of interest at a final concentration of 33 μM (which is in line with estimates of drug concentrations in the gastrointestinal tract)^23^, 2) a similar culture incubated with the same volume of a vehicle control (DMSO), and 3) a 3-ml volume of sterile mGAM, incubated with the same drug concentration. The no-drug control is important to distinguish microbiome-derived small molecules from ones that result from MDM of the tested drug. The no-microbiome control is important to distinguish cases of passive drug degradation or faulty chemical extraction from those of active MDM. Cultures and controls are then incubated for an additional 24 hours at 37°C in an anaerobic chamber, chemically extracted, and finally analyzed using High Performance Liquid Chromatography coupled with Mass Spectrometry (HPLC-MS). The entire procedure is repeated three consecutive times to verify the reproducibly of the screen.

To evaluate the feasibility, reproducibility, and scalability of our screen, we performed a pilot experiment on a selected set of 6 orally administered drugs that are diverse in structure and biological activities (erythromycin, antibiotic; terbinafine, antifugal; ketoprofen, antiinflammatory; valganciclovir, antiviral; topotecan, anticancer; atenolol, antihypertensive). Importantly, we also included a drug that is known to be readily metabolized by the human microbiome as a positive control: sulfasalazine, a prodrug that is intestinally activated by the human microbiome to produce the anti-inflammatory drug 5-aminosalicylic acid (5-ASA) and the metabolite sulfapyridine^24,25^. Unequivocally, we observed a reproducible metabolism of sulfasalazine into sulfapyridine, while the rest of the tested drugs remained unchanged in all three trials (**Fig. 2c, 2d**). These results establish our analytical screen as a valid method for determining the effect of MDM on orally administered drugs, where positive and negative results can be readily and reproducibly differentiated.

With these promising results from the pilot assay, we decided to apply MDM-Screen to a library of 575 orally administered drugs. This library is a subset of the SCREEN-WELL**^®^** FDA approved drug library (Enzo Life Sciences, Inc.), including only drugs with an established oral route of administration. We chose this library because of its diversity in chemical structure and pharmacological activity (**Supplementary Table 1**); and although all of the drugs in this library are currently being used in the clinic, almost nothing is known about their metabolism by the human gut microbiome. Following the procedures established in the pilot screen, we tested each drug twice, along with matching no-drug and no-microbiome controls. For final verification and consensus determination, a third trial was performed for drugs that showed a positive MDM on either or both of the first two trials. Therefore, a drug is deemed MDM+ when it is metabolized in the same manner during at least two out of three independent experiments. Taken together, we have developed and performed a high-throughput screen for mapping the ability of the complex human microbiome to metabolize orally administered small molecule drugs, in a systematic and unbiased manner.

### MDM-Screen identifies novel drug-microbiome interactions

Among the 575 drugs tested, 438 (76%) of them were successfully analyzed using our aforementioned procedures; the remaining 137 failed MDM-Screen due to issues related to drug stability or incompatibilities with the extraction or chromatography methods employed (see **Discussion**). Among the successfully analyzed drugs, 57 (13%) were identified as MDM-Positive (MDM+) (**Supplementary Table 1, Supplementary Table 2, and Supplementary Fig. 3**). As expected, several previously reported MDM cases were identified, further verifying MDM-Screen as a systematic method for discovering microbiome-drug interactions. These include the nitroreduction of the muscle relaxant dantrolene^26^, nitroreduction of the antiepileptic clonazepam (reported only in rats before this study)^27^, hydrolysis of the isoxazole moiety in the antipsychotic risperidone^28,29^, as well as several modifications to the bile acids chenodeoxycholic acid and ursodiol^30^.

**Figure 3.**
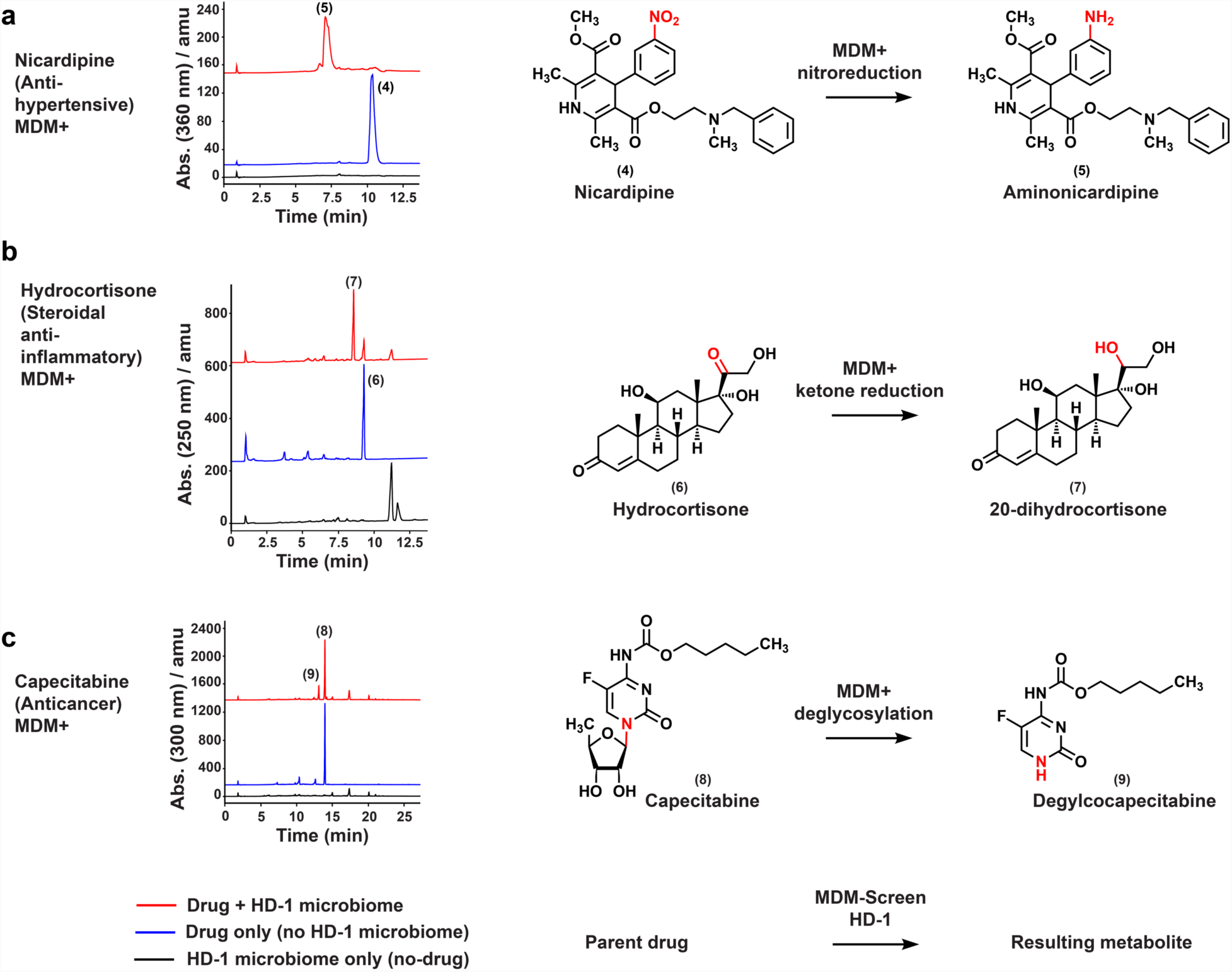
Examples of positive hits from MDM-Screen. An HPLC-MS analysis is shown for each of the selected examples, where three chromatograms are displayed per case: one for the drug incubated with HD-1 mGAM.02 culture (red), a second one for the drug incubated with mGAM.02 broth (blue), and a third one for HD-1 mGAM.02 culture with no drug added (black). **a)** An HPLC chromatogram at an absorbance of 360 nm is shown for nicardipine (4), indicating its conversion to aminonicardipine (5) in the presence of the HD-1 microbiome. **b)** An HPLC chromatogram at an absorbance of 250 nm is shown for hydrocortisone (6), indicating its conversion to 20-dihydrocortisone (7) in the presence of the HD-1 microbiome. **c)** An HPLC chromatogram at an absorbance of 300 nm is shown for capecitabine (8), indicating its conversion to deglycocapecitabine (9) in the presence of the HD-1 microbiome. Structures of the three metabolites were elucidated using NMR (**see Supplementary Data 1**).

More importantly, MDM-Screen identified a suite of novel MDM cases (46 cases, 80% of the MDM+ drugs). Among those, we selected four examples for detailed characterization: the commonly used anti-hypertensive / cardiac drug nicardipine, the chemotherapeutic agent capecitabine, and finally, the two steroidal anti-inflammatory drugs hydrocortisone (cortisol) and hydrocortisone acetate (often administered rectally), which produce an identical MDM metabolite. To unequivocally determine the structure of the resulting metabolite for each of these cases, we scaled up the biochemical incubation with HD-1, isolated and purified each of the resulting metabolites, and elucidated their structures using Nuclear Magnetic Resonance (NMR) (**see Methods and Supplementary Data 1**). Nicardipine metabolite (aminonicardipine) corresponds to the nitroreduced form of the drug: a common modification by members of the gut microbiome but one that has not been reported for this drug (**Fig. 3a**). For hydrocortisone, we determined that MDM results in the reduction of the ketone group at C20, producing 20-dihydrocortisone (**Fig. 3b**). For hydrocortisone acetate, the same modification occurs but is accompanied with deacetylation of the C21 hydroxyl group **(Supplementary Fig. 3)**. While C20 reduction was previously reported for hydrocortisone^31,32^, neither deacetylation nor C20 reduction were reported for hydrocortisone acetate. For capecitabine, we show that MDM results in complete deglycosylation, again, a modification never reported for this drug (**Fig. 3c**). Taken together, these results establish MDM-Screen as a viable method for identifying both known and novel biochemical modifications of structurally and pharmacologically diverse drugs by the gut microbiome.

### A global analysis of MDM by HD-1

Other than discovering novel drug-microbiome interactions, the results of our systematic screen allow for an unbiased, global analysis of MDM. Overall, the 57 MDM+ drugs belonged to 28 pharmacological classes and an even more diverse set of structural classes (**Fig. 4a and Supplementary Table 2**). We hypothesized that members of the microbiome would be more likely to metabolize natural or naturally-derived compounds due to a higher probability of prior exposure. To test this hypothesis, we first annotated each of the MDM+ or MDM-drugs to one of three categories: naturally occurring molecules (i.e., molecules directly derived from humans, plants, or microbes; an example of this category is hydrocortisone; N=30), derivatives of naturally occurring molecules (i.e., a semisynthetic derivative or a close structural mimic of a natural product, an example of this category is hydrocortisone acetate; N=90), and synthetic molecules (an example of this category is nicardipine; N=318). Interestingly, by comparing the fraction of MDM+ drugs in the first two categories (natural + derivative, 26 out of 120, 21.6%) to that of the third category (synthetic, 31 out of 318, 10%), we revealed a significant difference (*p* < 0.001, two-tailed proportions z-test). Intrigued, we decided to examine differences in MDM at lower levels of drug classification. We observed a significantly higher hit rate among steroids (steroids: 14 out of 26, 53.8%; non-steroid: 43 out of 412, 10.4%, *p* < 0.001, two-tailed proportions z-test), including hormonal steroids, corticosteroids, bile acids, and derivatives thereof. In fact, the high hit rate of the steroid class is the major contributor to the observed difference between the hit rates of natural/derivative and synthetic groups, which is abolished upon exclusion of the steroids (non-steroid natural/derivative: 12 out of 96, 12.5%; non-steroid synthetic: 31 out of 316, 10%). The high hit rate among steroids is in-line with the idea that the microbiome is more likely to metabolize compounds it frequently encounters, as steroids (e.g., bile acids) are normally present in the gut, and at high concentrations^33^. The fact that ∼10% of fully synthetic molecules are metabolized by HD-1 indicates the presence of a yet-unexplored range of biochemical activities that are encoded by the gut microbiome, and are capable of recognizing foreign substrates.

**Figure 4.**
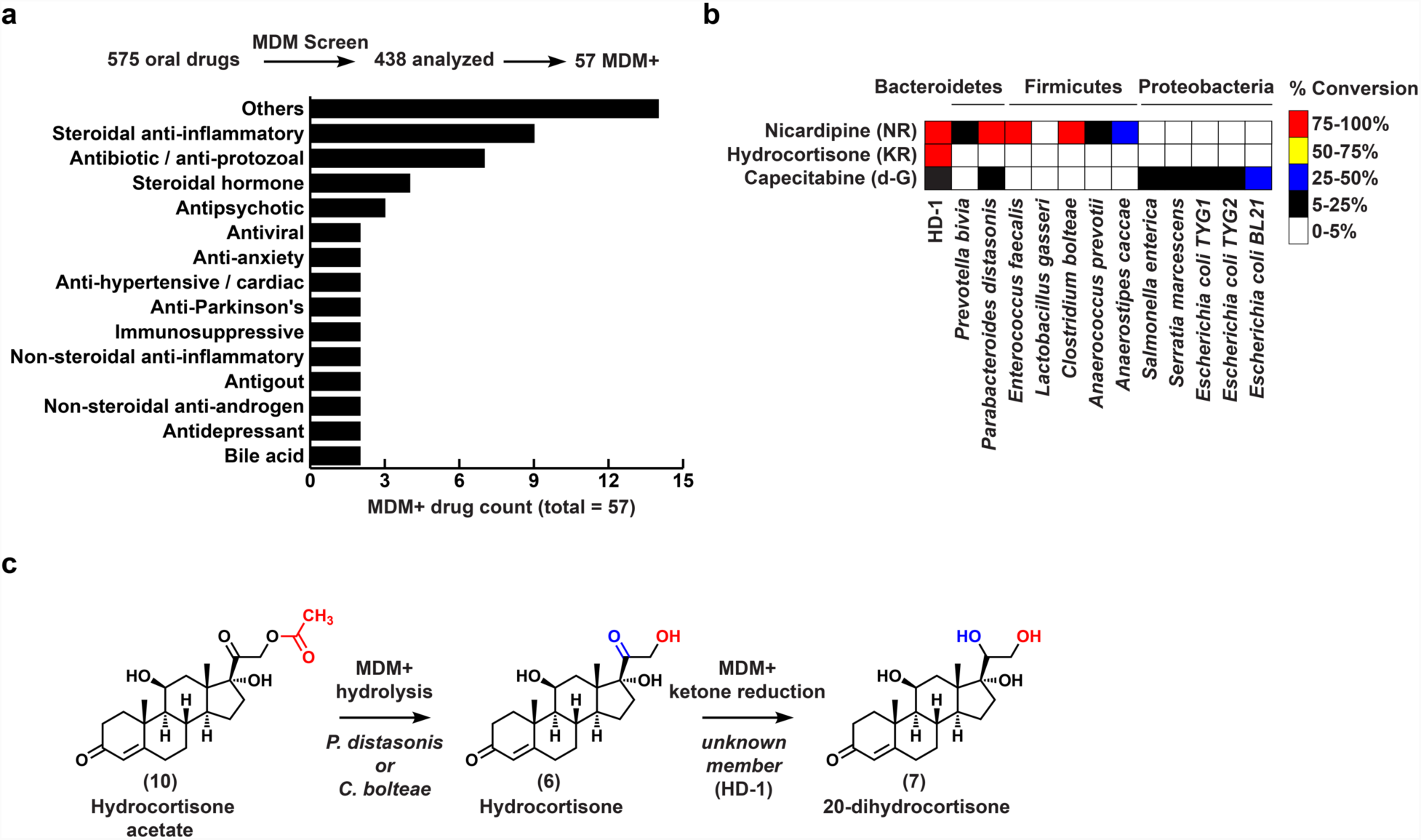
Overall results of MDM-Screen. **a)** A bar graph showing the pharmacological classes of MDM+ drugs discovered by MDM-Screen. “Others” include one drug each from 14 additional classes (**Supplementary Table 1**). In the inset bar graph, “natural”, “derivative”, or “synthetic” indicate whether the tested drug is: a natural product of any source (human, plant, microbial), a derivative of a naturally occurring molecule, or fully synthetic, respectively. *** indicates *p* < 0.001, two-tailed proportions z-test. **b)** A heat map indicating the ability of each of 12 tested strains to perform the three example modifications in **Fig. 3**: NR, nitroreduction; KR, ketone reduction; and d-G, deglycosylation. **c)** An example of sequential metabolism revealed by MDM-Screen: hydrocortisone acetate (10) can be first deacetylated by two members of the microbiome (*P. distasonis* and *C. bolteae*) to yield hydrocortisone (6), which is then reduced at C20 by a yet-unidentified member of the HD-1 microbiome to yield 20-dihydrocortisone (7). Structures were confirmed by NMR and comparison to authentic standards (**Supplementary Data 1 and Supplementary Fig. 4**).

### Linking MDM to specific members of the human microbiome

Our results from MDM-Screen indicate a significant and diverse ability of the collective gut microbiome to metabolize clinically used drugs that are unrelated in structure and biological activity. Next, we wondered whether the observed biochemical modifications can be attributed to specific members of the microbiome. To answer this question, we picked the same representative set of MDM transformations that we characterized above (3 transformations on 3 drugs) (**Fig. 3**), and explored the ability of a limited panel of 11 gut microbiome isolates and a laboratory strain to perform them. This panel was selected from three of the most abundant Phyla that normally inhabit the gut microbiome (Firmicutes, Bacteroidetes, and Proteobacteria), and spans 10 bacterial genera. Overall, nitroreduction of nicardipine was extensively performed by Bacteroidetes and Firmicutes, while capecitabine deglycosylation was mainly performed by Proteobacteria and one of the two tested Bacteroidetes: *Parabacteroides distasonis*. (**Fig. 4b**). None of the tested strains performed C20 reduction of hydrocortisone, suggesting that it is performed by a yet unidentified member(s) of the HD-1 microbiome (only two gut isolates were previously shown to perform C20 reduction on hydrocortisone: *Clostridium scindens* and *Butyricicoccus desmolans*)^31,32^.

Interestingly, we also observed sequential MDM transformations that appear to be contributed by different members of the microbiome on the same parent drug. An example of this includes the deacetylation (ester hydrolysis) and further reduction of hydrocortisone acetate. When hydrocortisone acetate (10) is incubated with either *P. distasonis* or *Clostridium bolteae*, it is deacetylated to yield hydrocortisone. When incubated with HD-1, however, it is both deacetylated and further reduced to yield 20-dihydrocortisone (**Fig. 4c and Supplementary Fig. 4**). Since we determined that a yet-unidentified member of the HD-1 microbiome is able to reduce hydrocortisone (6) at C20 (**Fig. 3 and Fig. 4b**), a two-step metabolic sequence is likely at play here, where hydrocortisone acetate (10) is first deacetylated to yield hydrocortisone (6) by *Parabcteroides* or *Clostridium* sp. in HD-1, then ketone reduced at C20 by another member of the microbiome to yield 20-dihydrocortisone (7). Overall, these results highlight the utility of our approach in mapping the ability of the complex human microbiome to metabolize drugs, whether it is contributed by one or several members of the microbiome: a key advance over experiments that are based on a single isolate.

### An MDM case study: capecitabine

Our ability to map MDM in a systematic manner is only the first step towards understanding the mechanistic details and biological consequences of direct drug-microbiome interactions. Therefore, we selected one MDM example, deglycosylation of capecitabine, for follow-up studies. Five main reasons motivated us to choose this modification for additional studies. First, the modification exerted on capecitabine yields a novel metabolite (deglycocapecitabine) that has not been previously reported in humans or animals, potentially providing more insights into the complex pharmacokinetics of this drug. Second, capecitabine is one of several generations of antimetabolite chemotherapeutic agents, many of which are prodrugs for 5-fluorouracil (5-FU), and are known collectively as the oral fluoropyrimidines (FPs)^34,35^. Because these agents share the same overall structure (a glycosylated and fluorinated pyrimidine), they may be subject to the same MDM. Third, oral FPs’ bioavailability and toxicity vary widely among patients^36,37^, but the human gut microbiome’s contribution to this variability has not been explored. Fourth, a related transformation was previously reported for another pyrimidine analog, the antiviral sorivudine, and linked to toxic outcomes during co-administration with 5-FU, suggesting the potential yet unexplored importance of deglycosylation for a wide range of drugs^38^. Finally, as shown above, capecitabine MDM is performed mainly by proteobacterial members of the microbiome, as well as some members of the Bacteroidetes. This feature not only provides genetically tractable organisms for functional studies (e.g., *E. coli*), but may also result in MDM variability between individuals depending on the relative abundance of specific metabolizers.

### Genetic basis of MDM deglycosylation

To gain more insights into the molecular mechanism of MDM deglycosylation, we sought to identify microbiome-derived enzymes responsible for this transformation. In humans, thymidine phosphorylase (TP) and uridine phosphorylase (UP), both part of the pyrimidine salvage pathway, were shown to catalyze the required deglycosylation of 5’-deoxy-5-fluorouridine at the last step of capecitabine metabolism to yield 5-FU^39^. To test whether bacterial homologs of human TP and/or UP are responsible for the observed MDM deglycosylation of capecitabine, we generated strains of *E. coli* BW25113 that are knockouts for TP (Δ*deoA*), UP (Δ*udp*), or both, and compared their ability to metabolize capectitabine to that of wild type *E. coli* (**Fig. 5a**). While wild type *E. coli* efficiently deglycosylates capecitabine (∼30% conversion rate), the deglycosylating activity of Δ*udp* and the Δ*deoA*/Δ*udp* knockout strains is significantly diminished (less than 4% conversion rate, *p*-value <0.001, two-tailed t-test) (**Fig. 5b**). Surprisingly, the Δ*deoA* knockout strain showed a significant increase in its deglycosylating activity in comparison to the wild type (∼ 50% conversion rate, *p*-value <0.01, two-tailed t-test), possibly due to a compensating mechanism (e.g., overexpression of *udp*) in the absence of *deoA*. These results indicate that microbiome-derived UP is, at least in part, responsible for the intestinal deglycosylation of capecitabine.

**Figure 5.**
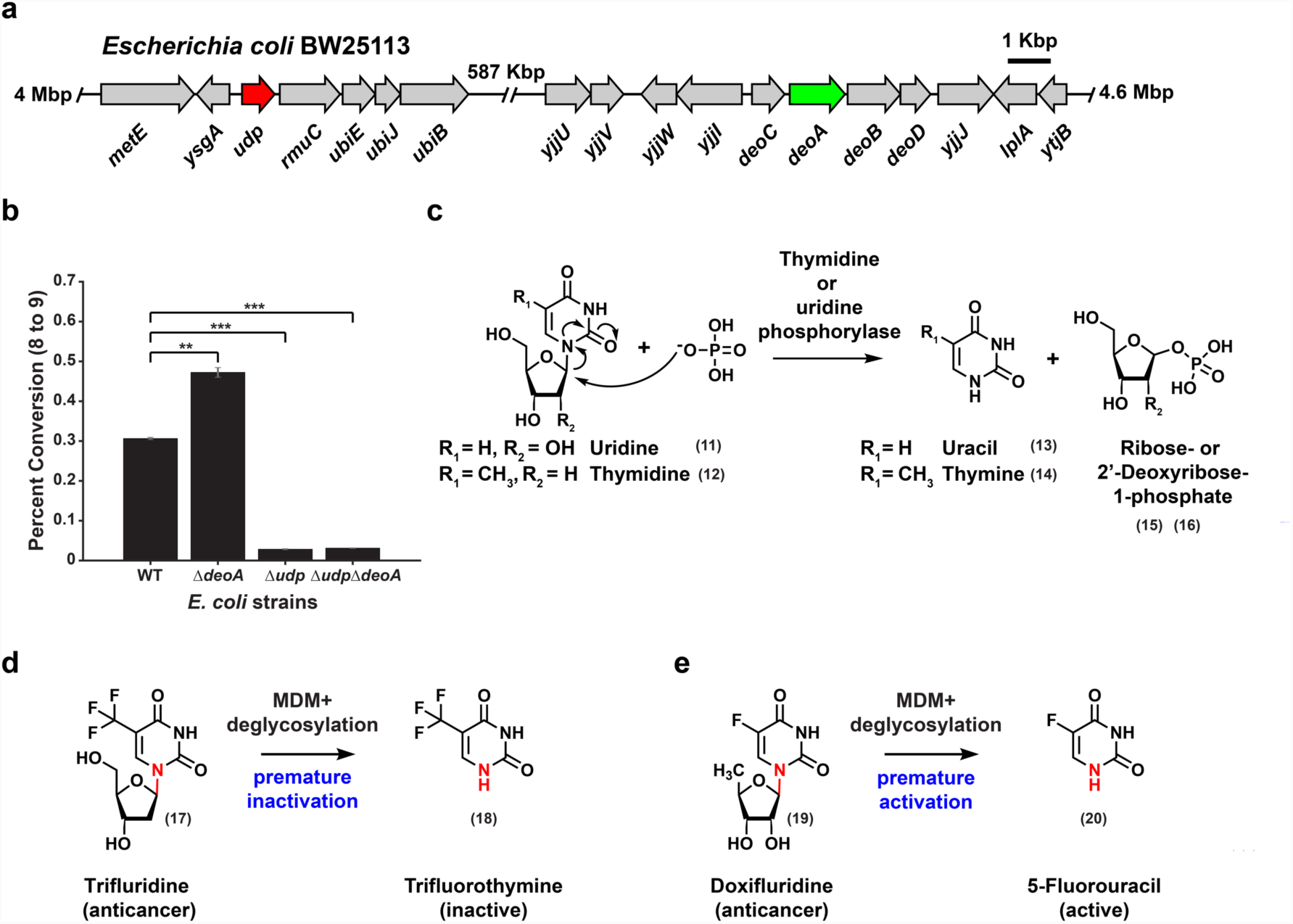
Genetic basis and widespread nature of MDM deglycosylation among the FPs. **a)** Genetic organization of the *udp* and *deoA* loci in the genome of *E. coli* BW25113. **b)** A bar graph indicating percent conversion of capecitabine (8) to deglycocapecitabine (9) by wild type *E. coli* BW25113 (WT), and Δ*udp*, Δ*deoA*, and Δ*deoA*/Δ*udp* mutants (each tested in triplicate). *** indicates *p*-value <0.001, while ** indicates *p*-value <0.01, two-tailed t-test. Error bars represent the standard deviation. **c)** Biochemical reaction catalyzed by thymidine and uridine phosphorylases on their natural substrates. **d)** MDM deglycosylation of the oral anticancer drug trifluridine (17) leads to its premature inactivation, since trifluorothymine (18) is no longer active. **e)** MDM deglycosylation of the anticancer prodrug doxifluridine (19) leads to its premature activation, since 5-fluorouracil (20) is the intended active metabolite. MDM deglycosylation of trifluridine and doxifluridine is also dependent on *deoA* and *udp*, and the structures of all resulting metabolites were confirmed by comparison to authentic standards (**Supplementary Fig. 5 and Supplementary Fig. 6**).

### MDM deglycosylation is widespread in the fluoropyrimidine class of chemotherapeutic agents

Next, we wondered whether deglycosylation occurs with other FPs, and whether the same enzymes are involved. To answer this question, we investigated the MDM of two additional oral FPs (doxifluridine and trifluridine), using both WT and mutant *E. coli*. We found that both drugs were subject to the same MDM deglycosylation, indicating that this modification is widespread among this class of molecules. Interestingly, unlike with capecitabine, almost complete deglycosylation was observed with WT *E. coli* (there was hardly any parent molecule left after 24 hours), and the activity was dependent on both TP and UP, as it was abolished only in the Δ*deoA*/Δ*udp* knockout (**Supplementary Fig. 5 and Supplementary Fig. 6**). These results indicate a level of deglycosylation specificity for TP/UP amongst the FPs, likely due to how well each drug mimics their natural substrate. Remarkably, the consequences of the same modification may be very different depending on the structural features of the tested drug. In the case of trifluridine, the resulting metabolite (trifluorothymine) is inactive (**Fig. 5d and Supplementary Fig. 5**): trifluridine needs to be incorporated intact into DNA to cause cytotoxicity^40^. Such a premature intestinal inactivation by the microbiome may thus be an unknown contributor to the established low bioavailability of trifluridine, in addition to the known contribution of human TP^36^. In the case of doxifluridine, however, the resulting metabolite is the active 5-FU itself (**Fig. 5e and Supplementary Fig. 6**). This premature activation of the prodrug may therefore lead into gastrointestinal toxicity – again, a side effect commonly associated with oral doxifluridine^41,42^.

**Figure 6.**
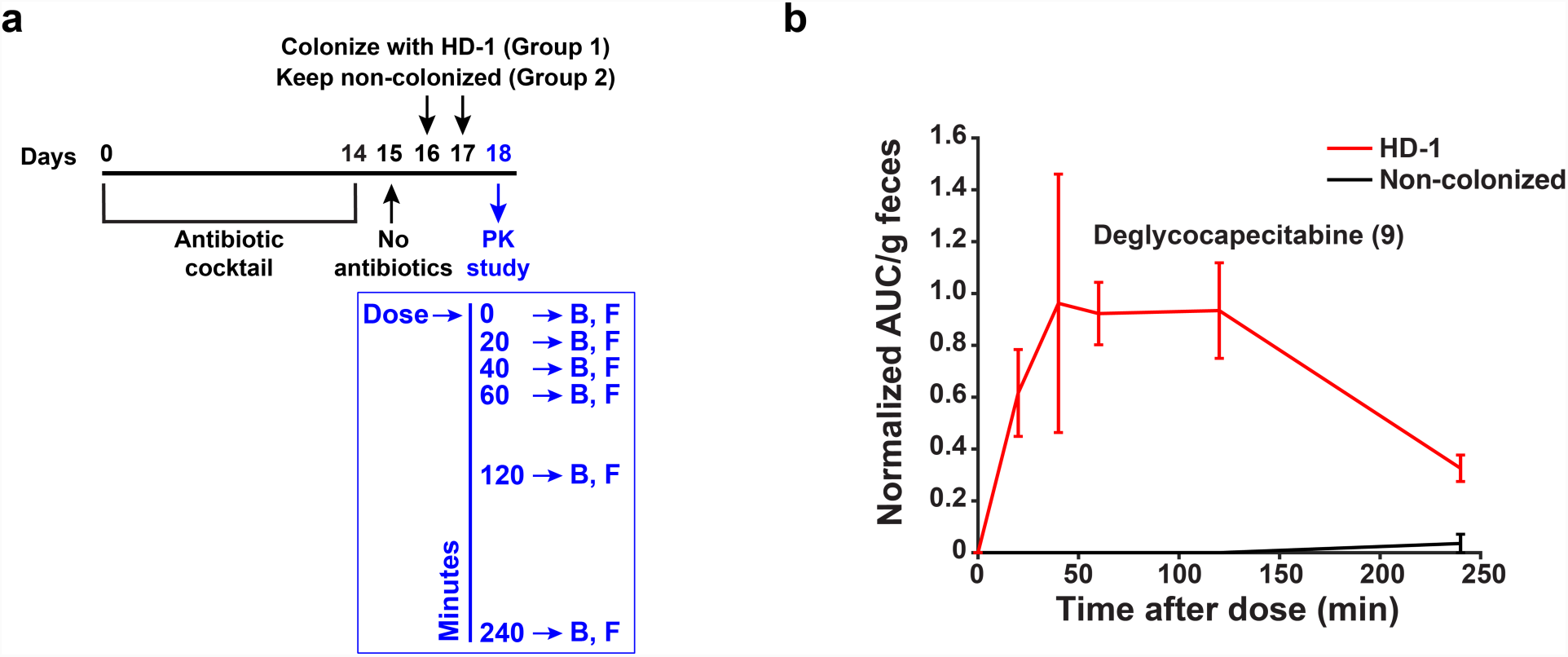
MDM deglycosylation occurs *in vivo*. **a)** Design of a microbiome-dependent pharmacokinetic experiment performed in mice using capecitabine. Mice are treated with antibiotics for 14 days, then colonized with HD-1 (N=6) or left non-colonized (N=6). On the pharmacokinetic experiment day, a single human-equivalent dose is administered to mice using oral gavage, and serial sampling of blood (B) and feces (F) is performed at 0, 20, 40, 60, 120, and 240 minutes post dosing. **b)** HR-HPLC-MS based quantification of deglycocapecitabine in fecal samples from mice colonized with HD-1 in comparison to non-colonized ones (see also **Supplementary Fig. 8**).Metabolite Area Under the Curve (AUC) per gram of feces is normalized by the AUC of an internal standard (voriconazole) (**see Methods**). Error bars represent the standard error of the mean.

To shed light on the potential consequences of capecitabine MDM deglycosylation, we sought to interrogate whether its metabolite, deglycocapecitabine, is able to re-enter the normal capecitabine metabolism cycle and yield 5-FU. In the liver, capecitabine is metabolized by liver carboxyesterases to yield 5’-deoxy-5-fluorocytidine, which is then deaminated by cytidine deaminase to yield 5’-deoxy-5-fluorouridine (doxifluridine). Preferentially in tumor tissues (due to the higher expression level of its metabolizing enzymes), doxifluridine is deglycosylated by human TP/UP to yield the active 5-FU (**Supplementary Fig. 7**)^43^. Similarly, deglycocapecitabine would almost certainly need to be processed by liver carboxyesterases to yield 5-fluorocytidine. Thus, we decided to directly test the activity of human carboxyesterase 1 (CES1) – the most important caroboxyesterase in capecitabine metabolism – against deglycocapecitabine^44,45^. Notably, while CES1 efficiently removed the carbamate group from capecitabine to yield 5’-deoxy-5-fluorocytidine *in vitro*, deglycocapecitabine was not recognized as a substrate by the enzyme under the same conditions (**Supplementary Fig. 7**). These results suggest that capecitabine MDM deglycosylation results in an inactivated product that is unlikely to yield the active 5-FU. Taken together, our findings indicate that FP deglycosylation is a common yet understudied MDM transformation that may have diverse consequences on the pharmacokinetics and/or pharmacodynamics of this widely used class of chemotherapeutic agents.

### MDM deglycosylation occurs *in vivo*

Although MDM-Screen was able to uncover novel microbiome-drug interactions, including MDM deglycosylation of FPs, it is unclear whether these results (observed *ex vivo*) can be recapitulated within a host (*in vivo*). To address this question, we selected MDM deglycosylation of capecitabine as a proxy for other FPs, and monitored it in an *in vivo* pharmacokinetic study that is performed in a microbiome-dependent manner. We treated two groups of C57/B6 mice with a cocktail of antibiotics for 14 days, then colonized one group with HD-1 while the control group remained non-colonized. The two groups were then treated with a single human-equivalent oral dose of capecitabine (755 mg/kg), and blood and feces were collected from each mouse at times 0, 20, 40, 60, 120 and 240 minutes post drug administration (**Fig. 6a**). Finally, we quantified capecitabine and its metabolites in chemical extracts from blood and feces using HR-HPLC-MS. Remarkably, deglycocapecitabine was detected in fecal samples from animals colonized with HD-1 as early as 20 min after dosing, and was almost completely absent in non-colonized ones (**Fig. 6b**). To our surprise, with the single dose regimen provided here, we could not detect deglycocapecitabine in mouse blood samples. In contrast, capecitabine, and its major liver-derived metabolite (5’-deoxy-5-fluorocytidine) were readily detected in blood with no significant differences between the two groups (**Supplementary Fig. 8**). These results indicate that – at least in the case of FP deglycosylation – MDM transformations observed *ex vivo* by MDM-Screen are recapitulated *in vivo*.

## DISCUSSION

In the current study, we develop a systematic screen for assessing the ability of the human gut microbiome to directly metabolize orally administered drugs, using a combination of microbial community cultivation, a high-throughput drug screen, bacterial genetics, and defined mouse colonization assays. Several key differences set our approach apart from previous studies in this area. First, instead of relying on single isolates in performing the initial screen, we use a well-characterized patient-derived microbial community that mimics to a large extent the original sample in composition and diversity. Despite the technical challenges associated with characterizing and maintaining stable microbial communities in batch cultures, three main advantages make this strategy worth pursuing: i) the extent of a biochemical transformation performed by single isolates cultured individually may be completely different than that performed by the same isolates when cultured as part of a complex community; ii) the net result of several members of the microbiome acting on the same drug can only be identified in mixed communities and not in single-isolate experiments, unless all pairwise and higher order permutations are tested; and iii) our strategy is “personalizable”. Some of the results obtained here – including the extent and type of certain modifications – will likely be specific to the strain-level composition of the HD-1 microbiome, and may vary if the assay is repeated with samples from different subjects. MDM-Screen has thus a good potential for assessing inter-patient variability in MDM.

Second, while most previous studies have focused on certain drug / species combinations that have historically been deemed important, our screen is agnostic towards the modifications being detected, the drugs being screened, and the responsible members of the microbiome being identified. This unbiased, systematic approach allowed us to map for the first time the potential extent of MDM, and to discover drug-microbiome interactions never reported before. We provide these results as a resource for the scientific community to further study the mechanistic details and pharmacological consequences of these newly discovered interactions. Third, MDM-Screen is performed in an efficient, high-throughput manner for both the organisms (a complex microbial community mimicking the original microbiome sample), and the drugs tested (almost 600 drugs were tested in an academic lab setting). With additional optimizations on the cultivation side (e.g., the use of 96-well plates) and the analytical chemistry side (e.g., automation of the extraction procedures), one can easily expand the screen to hundreds of human microbiome samples and thousands of drugs.

Despite these advances, our approach is still subject to several limitations. First, 24% of the drugs tested failed to be analyzed using the general analytical chemistry workflow described in the initial MDM-Screen. These drugs fell into one or more of three main categories: were not stable after overnight incubation in no-microbiome controls, could not be extracted using ethyl acetate, or could not be analyzed using reverse phase chromatography, with the last two being attributed mostly to polar or charged compounds. An alternative chemical analysis method will need to be developed for these molecules in order to assess their MDM. Second, we focused initially on oral drugs, yet several parenteral drugs and their liver-derived metabolites may be subject to important MDM transformations after biliary secretion. Third, even in our most diverse *ex vivo* cultures, we fail to support the growth of 100% of the community in the original sample. Finally, we initially based our analysis on a single human sample, HD-1. Therefore, it is almost certain that the types of MDM transformations observed here are an underestimation of all possible ones, and that performing MDM-Screen several times with samples derived from unrelated subjects may be necessary to reveal the complete biochemical potential of MDM.

Although MDM was shown to lead into changes in the bioavailability, toxicity, and/or efficacy of certain therapeutics (e.g., digoxin) – to the same extent as liver metabolism – it is almost entirely overlooked by the regulatory agencies when developing new drugs^14,46^. Our current study was designed to achieve two main goals: a) develop a simple platform for studying MDM in a systematic manner; b) map the extent of MDM against commonly used drugs, including the functional characterization of key proof-of-principle examples. By achieving these goals, our overall findings reveal an unexpectedly large and diverse ability of the human microbiome to directly metabolize clinically used, small molecule drugs, and a wide potential for MDM as a key factor in explaining the observed inter-patient variability in the pharmacokinetics and/or pharmacodynamics of these agents. At the same time, our approach provides the regulatory agencies (e.g., the Food and Drug Administration) with a simple screen for assessing MDM that can be easily implemented in any typical drug development pipeline. It is crucial that drug-microbiome interactions, including both effects of drugs on the microbiome (which were systematically mapped in an elegant screen published recently)^23^, as well as MDM (mapped here for the first time) are considered while studying the pharmacology and toxicology of newly developed therapeutic agents.

## METHODS

### *ex vivo* culture of human gut microbiome communities

The Institutional Review Board (IRB) at Princeton University determined that the activity was not human subjects research. Consequently, Princeton IRB approval was not applicable. Freshly collected human fecal material from a healthy donor, HD-1 (∼ 30 min from collection, transported on ice) was brought into an anaerobic chamber (70% N_2_, 25% CO_2_, 5% H_2_). One gram of the sample was suspended in 15 ml of sterile phosphate buffer (PBSc) supplemented with 0.1% L-cysteine in a 50 ml sterile falcon tube. The suspension was left standing still for 5 min to let insoluble particles settle. The supernatant was mixed with an equal volume of 40% glycerol in PBSc. Aliquots (1 ml) of this suspension were placed in sterile cryogenic vials and frozen at −80°C until use^47^.

A small aliquot (∼20 μl) from an HD-1 glycerol stock was used to inoculate 10 ml of 14 different media: Liver Broth (Liver), Brewer Thioglycolate Medium (BT), Bryant and Burkey Medium (BB), Cooked Meat Broth (Meat), Thioglycolate Broth (TB), Luria-Bertani Broth (LB) (obtained from Sigma Aldrich, USA), Brain Heart Infusion (BHI), MRS (MRS), Reinforced Clostridium Medium (RCM), M17 (M17) (obtained from Becton Dickinson, USA), modified Gifu Anaerobic Medium (mGAM) (obtained HyServe, Germany), Gut Microbiota Medium (GMM^47^), TYG, and a 1:1 mix of each (BestMix), and cultures were incubated at 37 °C in an anaerobic chamber. One ml was harvested from each culture each day for 4 consecutive days, and centrifuged to recover the resulting bacterial pellets. DNA was extracted from all pellets using the Power Soil DNA Isolation kit (Mo Bio Laboratories, USA), the 16S rRNA gene was amplified (∼250 bps, V4 region), and Illumina sequencing libraries were prepared from the amplicons according to a previously published protocol and primers^48^. Libraries were further pooled together at equal molar ratios and sequenced on an Illumina HiSeq 2500 Rapid Flowcell as paired-end (2X175 bps) reads, along with 8 bps Index reads, following the manufacturer’s protocol (Illumina, USA). Raw sequencing reads were filtered by Illumina HiSeq Control Software to generate Pass-Filter reads for further analysis. Different samples were de-multiplexed using the index reads. Amplicon sequencing variants (ASVs) were then inferred from the unmerged pair-end sequences using the DADA2 plugin within QIIME2 version 2018.6^16,17^. The forward reads were trimmed at 165 bp and the reverse reads were trimmed at 140 bp. All other settings within DADA2 were default. Taxonomy was assigned to the resulting ASVs with a naive Bayes classifier trained on the Greengenes database version 13.8^18,19^. Only the target region of the 16S rRNA gene was used to train the classifier. Rarefaction analysis was performed within QIIME2 ^17^.

### *ex vivo* screening of the drug library

In an anaerobic chamber, a small (∼100 μl) of an HD-1 glycerol stock was diluted in 1 ml of mGAM, then 20 μl of this solution was used to inoculate 3 ml of mGAM in culture tubes. Cultures were grown for 24 hours at 37 °C in an anaerobic chamber. After 24 hours, 10 μL of each drug (the concentration of each molecule in the library is 10 mM), or of a DMSO control were added to the growing microbial community. In addition, 10 μL of each drug was also incubated similarly in a no-microbiome, mGAM control. HD-1 / DMSO control pellets from several batches of the screen were analyzed using high-throughput 16S rRNA gene sequencing as described above to ensure the maintenance of a similarly diverse microbial composition. Experiments and controls were allowed to incubate under the same conditions for a second 24-hour period. After incubation, cultures were extracted with double volume of ethyl acetate and the organic phase was dried under vacuum using a rotary evaporator (Speed Vac). The dried extracts were suspended in 250 μL MeOH, centrifuged at 15000 rpm for 5 min to remove any particulates, and analyzed using HPLC-MS (Agilent Single Quad, column: Poroshell 120 EC-C18 2.7um 4.6 × 50mm, flow rate 0.8 ml/min, 0.1% formic acid in water (solvent A), 0.1% formic acid in acetonitrile (solvent B), gradient: 1 min, 0.5% B; 1-20 min, 0.5%-100% B; 20-25 min, 100% B). If drugs were deemed positive for MDM in one or both of the two runs, they were analyzed a third time using both HPLC-MS and HR-HPLC-MS/MS (Agilent QTOF, column: Poroshell 120 EC-C18 2.7um 2.1×100 mm, flow rate 0.25 ml/min, 0.1% formic acid in water (solvent A), 0.1% formic acid in acetonitrile (solvent B), gradient: 1 min, 0.5% B; 1-20 min, 0.5%-100% B; 25-30 min, 100% B). For selected molecules, cultures were scaled up and metabolites were purified and their structures were elucidated using NMR (see below).

### Isolation and structural elucidation of selected metabolites

1 ml of HD-1 glycerol stock was used to inoculate 100 ml mGAM medium and cultured for 24 hours at 37 °C in an anaerobic chamber. After 24 hours, 2 ml of 10 mM capecitabine, hydrocortisone or nicardapine solutions were added to the HD-1 culture and incubated for another 24 hours. After the second 24 hours, the cultures were extracted with double the volume of ethyl acetate and the organic solvent layer was dried under vacuum in a rotary evaporator. The dried extract was then suspended in MeOH and partitioned by reversed phase flash column chromatography (Mega Bond Elut-C18 10g, Agilent Technology, USA) using the following mobile phase conditions: solvent A, water with 0.01% formic acid; solvent B acetonitrile with 0.01% formic acid, gradient, 100% A to 100% B in 20% increments. Fractions containing the metabolites of interest were identified by HPLC-MS, and reverse phase HPLC was used to purify each metabolite using a fraction collector (Agilent Single Quad, column Poroshell 120 EC-C18 2.7 um 4.6×100 mm, flow rate 0.8 ml/min, 0.1% formic acid in water (solvent A), 0.1% formic acid in acetonitrile (solvent B), gradient: 1 min, 0.5% B; 1-30 min, 0.5%-100% B; 30-35 min, 100% B). The purified metabolites were subjected to NMR and HR-MS/MS analysis. Structural elucidation details of capecitabine, hydrocortisone, and nicardipine metabolites are detailed in **Supplementary Data 1**.

### MDM-Screen using a panel of representative isolates from the gut microbiome

3 ml of pre-reduced medium (PYG, RCM, GAM, BHI or LB, depending on the isolate, incubated for 24 hours in the anaerobic chamber) was inoculated with the corresponding isolate’s glycerol stock. Cultures were grown overnight at 37°C in an anaerobic chamber (70% N_2_, 25% CO_2_, 5% H_2_). 20 μL of these seed cultures were inoculated into 3 ml of the same selected medium, and incubated at 37°C under the same anaerobic conditions for an additional 24 hours. After 24 hours, 10 μL of the 10 mM drug solution in DMSO, or of a DMSO control were added to the growing microbial culture and incubated for another 24 hours. In addition, 10 μL of each drug were incubated for 24 hours under the same conditions in a no-bacterium, medium-only control. After incubation, cultures were extracted with ethyl acetate and the organic phase was dried under vacuum in a rotary evaporator. Extracts were suspended in 250 μl of MeOH and analyzed using HPLC-MS as described above.

### TP and UP gene deletions in *E. coli* BW25113

*E. coli* BW25113 mutants that harbor a replacement of *deoA* or *udp* with a kanamycin resistance gene were obtained from the Keio collection^49^. Since the kanamycin resistance gene is flanked by FLP recognition target sites, we decided to excise it and obtain in-frame deletion mutants. Plasmid pCP20, encoding the FLP recombinase, was transformed to each of the mutants by electroporation, and transformants were selected on Ampicillin at 30 °C for 16 hours. 10 transformants from each mutant were then picked in 10 μl LB medium with no selection, and incubated at 42 °C for 8 hours to cure them from the temperature-sensitive pCP20 plasmid. Each growing colony was then streaked on three plates (LB-ampicillin, LB-kanamycin, and LB with no selection). Mutants that could only grow on LB, but not on LB-ampicillin (confirming the loss of the pCP20 plasmid), nor on LB-kanamycin (confirming the excision of the kanamycin resistance gene) were confirmed to harbor the correct deletion using PCR and DNA sequencing. Primers deoA-Check-F: 5’-CGCATCCGGCAAAAGCCGCCTCATACTCTTTTCCTCGGGAGGTTACCTTG-3’, deoA-Check-R: 5’-CAAATTTAAATGATCAGATCAGTATACCGTTATTCGCTGATACGGCGATA-3’, udp-Check-F: 5’-CGCGTCGGCCTTCAGACAGGAGAAGAGAATTACAGCAGACGACGCGCCGC-3’, and udp-Check-R: 5’-TGTCTTTTTGCTTCTTCTGACTAAACCGATTCACAGAGGAGTTGTATATG-3’ were used in PCR experiments to confirm the deletion of the *deoA* or *upd* genes and the kanamycin resistance gene replacing them^49^. To construct the Δ*deoA*/Δ*udp* double knockout, the in-frame Δ*udp* knockout obtained above was used as a starting point. Plasmid pKD46 expressing the λ Red recombinase was transformed to it using electroporation,^50^ and transformants were selected on LB-Ampicillin at 30 °C for 16 hours. One Ampicillin-resistant transformant was then cultured at 30 °C in 50 ml of LB-Ampicillin, with an added 50 μl of 1 M L-arabinose to induce the expression of the recombinase. At an optical density of 0.4-0.6, electrocompetent cells were prepared from the growing culture by serial washes in ice cold 10% glycerol, and ∼300 ng of a linear PCR product were transformed to it by electroporation. This PCR product was prepared by using the deoA-Check-F and deoA-Check-R primers on a template DNA prepared from the *deoA* mutant of the Keio library, in which a kanamycin resistance gene replaces *deoA*. After electroporation, transformants were selected on LB-kanamycin at 37 °C to induce the loss of the temperature sensitive pKD46 plasmid, cultured in LB-kanamycin overnight at 37 °C, and checked by PCR to confirm the correct recombination position. Finally, the kanamycin resistance gene was excised from the *deoA* locus by the FLP recombinase using the same strategy explained above, resulting in the final Δ*deoA*/Δ*udp* mutant.

### MDM-Screen of capecitabine using wild type and mutant *E. coli*

Wild type *E. coli* BW25113, and corresponding TP knockout (Δ*deoA*), UP knockout (Δ*udp*), and TP/UP double knockout (Δ*deoA*/Δ*udp*) strains were cultured overnight in LB medium (aerobically, shaking at 37 °C, 50 ml each). Triplicates of 3 ml for each strain were incubated with 10 μl of 10 mM capecitabine (in DMSO) for an additional 24 hours in an anaerobic chamber along with bacteria-only and media-only controls. Cultures were then extracted and analyzed as previously described, except for the addition of 20 μL of 0.25 mg/ml of an internal standard (voriconazole) prior to the extraction.

### MDM-Screen of other FPs using wild type and mutant *E. coli*

Wild type *E. coli* BW25113, and corresponding TP knockout (Δ*deoA*), UP knockout (Δ*udp*), and TP/UP double knockout (Δ*deoA*/Δ*udp*) strains were cultured overnight in LB medium (aerobically, shaking at 37 °C, 50 ml each). Aliquots (100 μl) of each strain were used to inoculate 3 ml of M9 medium, which were grown again overnight (aerobically, shaking at 37 °C). 10 μl of 10 mM doxifluridine (in DMSO) or trifluridine (in methanol) were incubated with each culture for an additional 24 hours in an anaerobic chamber, along with bacteria-only and medium-only controls. Cultures were spun down and collected supernatants were lyophilized. The dried residues were then resuspended in 500 μL methanol and analyzed by HPLC-MS (Agilent Single Quad; column: Poroshell 120 EC-C18 2.7um 4.6 × 100mm; flow rate: 0.6 ml/min; solvent A: 0.1% formic acid in water: solvent B: 0.1% formic acid in acetonitrile) and the following gradient: 1 min, 0.5% B; 1-20 min, 0.5%-35% B; 25-30 min, 35%-100% B; 30-35 min, 100% B.

### Microbiome-dependent pharmacokinetic experiment

All animal experiments were conducted according to USA Public Health Service Policy of Humane Care and Use of Laboratory Animals. All protocols were approved by the Institutional Animal Care and Use Committee, protocol 2087-16 (Princeton University). 8-10-weeks old (25-30 g) C57BL/6 mice were purchased from Jackson laboratories. 12 mice were treated with a commonly used cocktail of antibiotics (1 g/l of amplicilin, neomycin, metronidazole and 0.5 g/l vancomycin) in drinking water for 14 days^51^. The antibiotic solution was supplemented with 5 g/l aspartame to make it more palatable^52^. During these two weeks, the gut microbiome composition was monitored by collecting feces from each mouse and performing molecular and microbiological analyses to make sure the microbiome is being cleared by the antibiotic treatment. On day 15, no antibiotics are administered for 24 hours (a washout period). On day 16, mice were separated into the two groups, 6 per group (3 males and 3 females). In group 1, mice remained non-colonized. In group 2, mice were administered 200 μl of freshly thawed HD-1 glycerol stock using oral gavage. On day 17, the oral gavage was repeated the same way to ensure the colonization of the administered bacteria (fecal samples were collected on days 16 and 17 and cultured anaerobically to ensure colonization). On day 18, the pharmacokinetic experiment was performed by monitoring the fate of capecitabine in mouse blood and feces over time. A capecitabine dose equivalent to a single human dose and adjusted to the weight of the mice was administered by oral gavage (755 mg / kg, as a solution in 50 μl DMSO), then serial sampling of tail vein blood (by tail snipping), as well as fecal collection were performed at these time points (zero, 20 min, 40 min, 60 min, 2 hours, and 4 hours). Blood for each time point (30 μl) was collected using a 30 μl capillary tube and bulb dispenser (Drummond Microcaps, Drummond Scientific), quickly dispensed in 60 μl EDTA to prevent blood coagulation, and stored on ice for up to 4 hours and then frozen at −80 °C until further analysis. Feces were also collected at the same time points (even though defecation was left at will, we succeeded in collecting feces for most time points), stored on ice for up to 4 hours and then frozen in −80 °C until further analysis. After the 4-hour pharmacokinetic time point, mice were euthanized.

For chemical extraction, 2 μl of an internal standard solution (0.5 mg / ml of voriconazole) were added to the blood / EDTA solution mentioned above, and the sample was mixed using a vortex mixer. Next, 500 μl of ethyl acetate was added and mixed. The sample was then centrifuged briefly at 15000 rpm, and the organic layer was transferred to a glass tube and evaporated under vacuum using rotary evaporation (Speed Vac). The dried residue was dissolved in 100 μl of MeOH, and the solution was centrifuged at 15000 rpm and transferred to an autosampler vial for HR-HPLC-MS analysis. For fecal samples, pellets were weighed (for later normalizations), and suspended in 500 μl sterile Milli-Q water (Millipore Corporation, USA). 2 μl of an internal standard solution (0.5 mg / ml of voriconazole) were added to the sample, and the mixtures were extracted with 500 μl 1:1 ethyl acetate : MeOH. Fecal debris were then spun down and collected supernatants were dried under vacuum using a rotary evaporator (Speed Vac). The dried residues were suspended in 100 μl MeOH. The final solutions were centrifuged at 15000 rpm and transferred to autosampler vials.

The prepared samples were analyzed by HR-HPLC-MS (Agilent QTOF).Chromatography separation was carried out on a Poroshell 120 EC-C18 2.7 um 2.1 × 100 mm column (Agilent, USA) with the gradient: 99.5% A, 0.5% B to 100% B in 20 minutes and a flow rate of 0.25 ml/min, where A= 0.1% formic acid in water and B= 0.1% formic acid in acetonitrile. A 10 μl aliquot of the reconstituted extract was injected into the HR-HPLC-MS system, and the Area Under the Curve (AUC) was integrated for each metabolite and normalized by the internal standard’s AUC. Peak identities were confirmed by accurate mass, and by comparison of chromatographic retention time and MS/MS spectra to those of authentic standards.

### *in vitro* metabolism of capecitabine and deglycocapecitabine using human carboxyesterase 1

Human Carboxylesterase 1 (CES1) was purchased from Sigma-Aldrich. Capecitabine or deglycocapecitabine (3.25 μl of a 10 mM stock in DMSO) was added into CES (50 μg) in 20 mM HEPES pH 7.4; in a total volume of 150 μL and then incubated at 37 °C^53^. After 60 min, the reaction was quenched with 150 μL of acetonitrile and placed on ice. The mixture was centrifuged for 3 min at 15000 rpm. The supernatant was dried under vacuum using rotary evaporation (Speed Vac). The dried residues were suspended in 200 μL MeOH. The final solutions were centrifuged at 15000 rpm and transferred to autosampler vials. The samples were analyzed by HPLC-MS (Agilent Single Quad 6120) for metabolite formation (column: Poroshell 120 EC-C18 2.7um 4.6 × 50mm, flow rate 0.8 ml/min, 0.1% formic acid in water (solvent A), 0.1% formic acid in acetonitrile (solvent B), gradient: 1 min, 0.5% B; 1-30 min, 0.5%-100% B; 30-35 min, 100% B).

## Data availability

All data reported in this study are included in this manuscript and accompanying Supplementary Information.

## Acknowledgments

We would like to thank Wei Wang and the Lewis Sigler Institute sequencing core facility for assistance with high-throughput 16S rRNA gene amplicon sequencing, Matthew Cahn for assistance with sequencing data analysis, Joseph Koos, A. James Link, and Yuki Sugimoto for assistance with Mass Spectrometry, Riley Skeen-Gaar for assistance with statistical analysis, Joseph Sheehan and Zemer Gitai for assistance with obtaining the Keio library mutants, Laboratory Animal Resources at Princeton University for assistance with mouse studies, and members of the Donia lab for useful discussions. Funding for this project has been provided by an Innovation Award from the Department of Molecular Biology, Princeton University, and an NIH Director’s New Innovator Award (ID: 1DP2AI124441), both to M.S.D. B.J. is funded by a New Jersey Commission on Cancer Research Pre-doctoral award (ID: DFHS18PPC056), and J.L is funded by a National Science Foundation Graduate Research Fellowship (ID: 2017249408).

## Author contributions

M.S.D., P.C., and B.J. designed the study. P.C., B.J., J.L., R.H., S.C. and M.S.D. performed experiments and analyzed the data. M.S.D., P.C., B.J., and J.L. wrote the manuscript.

## Supplementary Information

Supplementary Tables, Supplementary Figures, and Supplementary Data are provided.

