## Supplementary Figures for "Systematic mapping of drug metabolism by the human gut microbiome"

**Contents:**

**Supplementary Figure 1.** A four-day time course of HD-1 cultured *ex vivo* in various media.

**Supplementary Figure 2.** Rarefaction analysis of HD-1 *ex vivo* cultures.

**Supplementary Figure 3.** Chromatograms of MDM+ Drugs.

**Supplementary Figure 4.** Sequential metabolism of hydrocortisone acetate by the HD-1 microbiome.

**Supplementary Figure 5.** MDM deglycosylation of the anticancer drug trifluridine.

**Supplementary Figure 6.** MDM deglycosylation of the anticancer prodrug doxifluridine.

**Supplementary Figure 7.** Capecitabine and deglycocapecitabine metabolism by human and bacterial enzymes.

**Supplementary Figure 8.** Microbiome-dependent pharmacokinetics of capecitabine.

**Other supplementary information not included in this file:**

**Supplementary Table 1.** Overall MDM-Screen results for the entire SCREEN-WELL® FDA approved drug library.

**Supplementary Table 2.** Detailed analysis of MDM+ Drugs.

**Supplementary Data 1.** Structural elucidation of selected MDM metabolites

**Supplementary Figure 1. A four-day time course of HD-1 cultured *ex vivo* in various media.** Family level bacterial composition of the original HD-1 fecal sample (far left), as well as that of HD-1 *ex vivo* cultures grown anaerobically in 14 different media over four days (.01, .02, .03, .04). 16S rRNA gene sequences that could not be classified at the family level, and families with less than 1% relative abundance in all samples are grouped into “Other”. Cultures are ordered according to their Jensen-Shannon ($D_{JS}$) divergence from the original HD-1 sample (upper axes, computed at the family level), where lower values indicate higher similarity to HD-1. Note that cultures grown in mGAM are the most similar to HD-1.

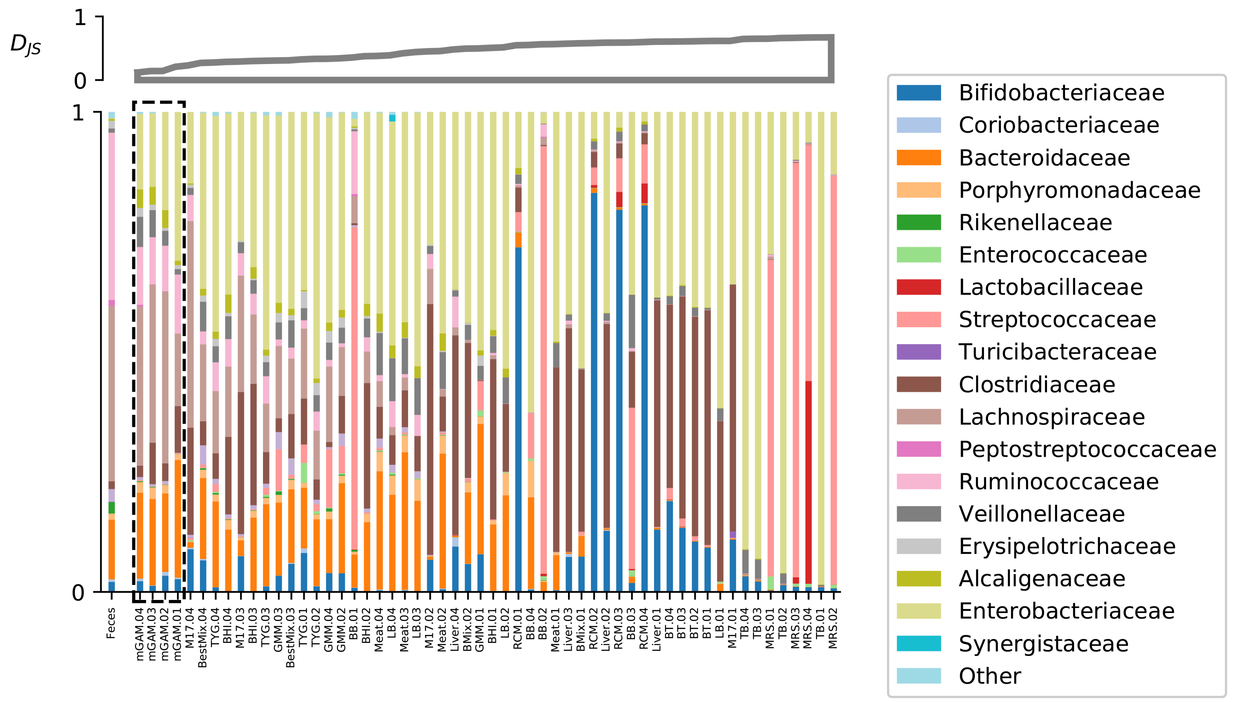

**Supplementary Figure 2. Rarefaction analysis of HD-1 *ex vivo* cultures.** Shannon diversity rarefaction of the first two days of each culture condition and the original HD-1 sample. Shannon diversity is computed at the ASV level. To better show the rise of the curve, only the first 40,000 sequences are shown out of ~100,000 per sample. The results do not change appreciably with the inclusion of the remaining sequences.

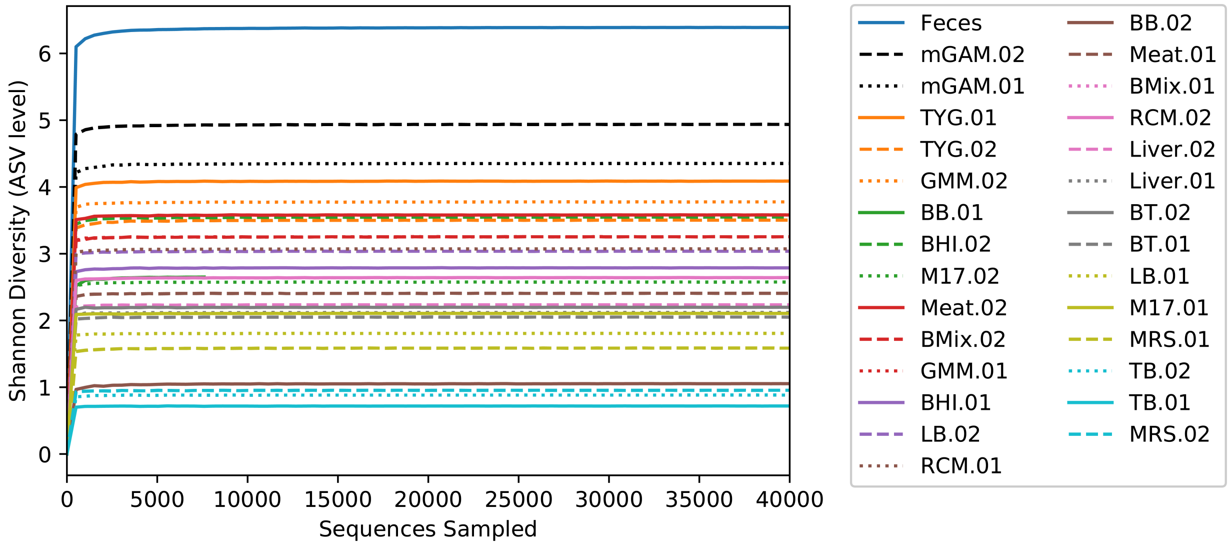

**Supplementary Figure 3. Chromatograms of MDM+ Drugs. a-f)** HPLC-MS analyses for each of the identified MDM+ drugs, where three chromatograms are displayed per case: one for the drug incubated with HD-1 mGAM.02 culture (red), a second one for the drug incubated with mGAM.02 broth (blue), and a third one for HD-1 mGAM.02 culture with no drug added (black). “p” indicates parent drug, “m1, m2, etc.” indicate the identified metabolites. HPLC chromatograms at an indicated wavelength, or MS Total Ion Chromatograms, are shown for each case. The name of the analyzed drug and the library well position are indicated above. See **Supplementary Tables 1 and 2** for additional information about the drugs and their identified metabolites.

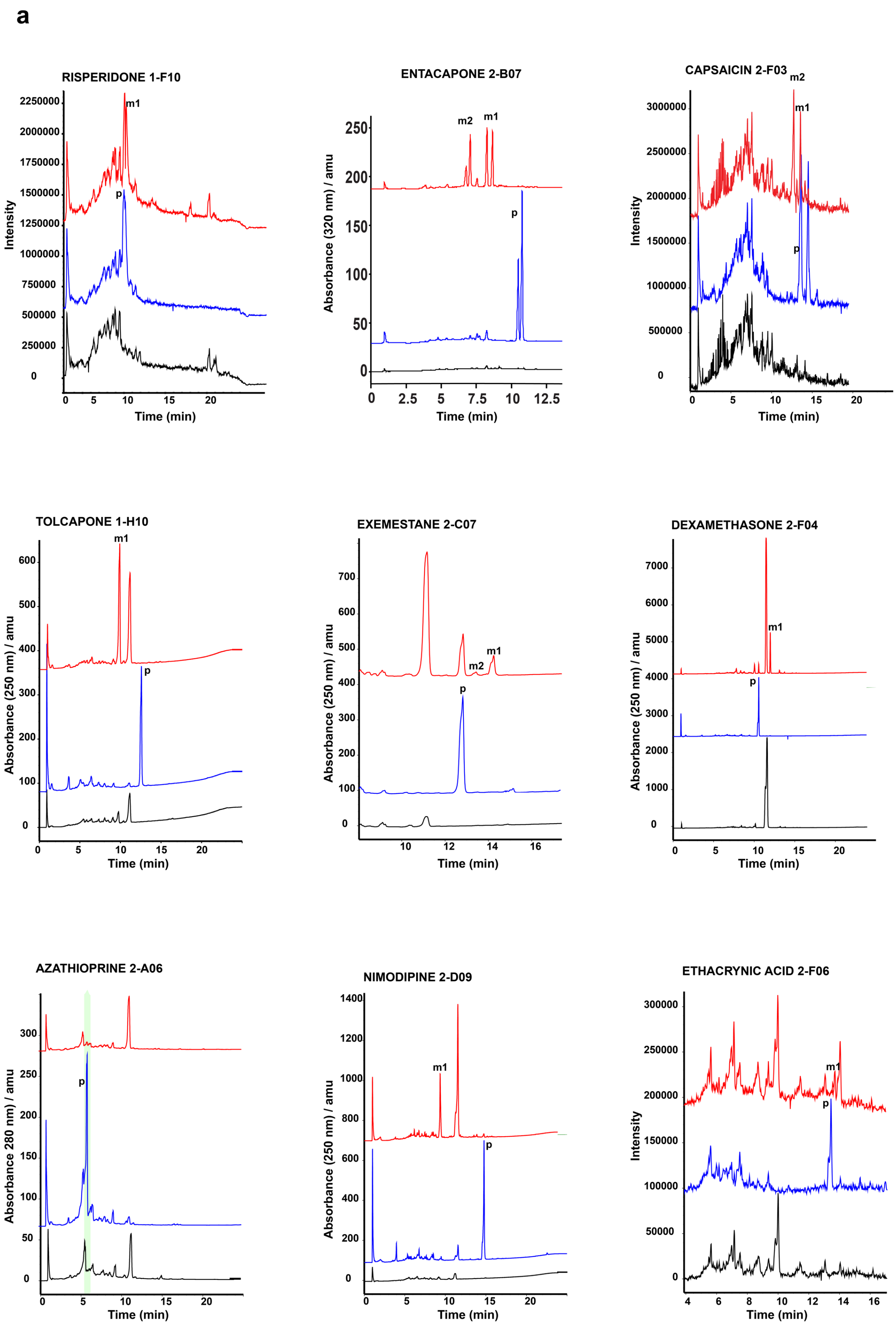

a
RISPERIDONE 1-F10
ENTACAPONE 2-B07
CAPSAICIN 2-F03
TOLCAPONE 1-H10
EXEMESTANE 2-C07
DEXAMETHASONE 2-F04
AZATHIOPRINE 2-A06
NIMODIPINE 2-D09
ETHACRYNIC ACID 2-F06
Intensity
Absorbance (320 nm) / amu
Absorbance (250 nm) / amu
Absorbance 280 nm) / amu
Time (min)
m1
m2
p

**b**

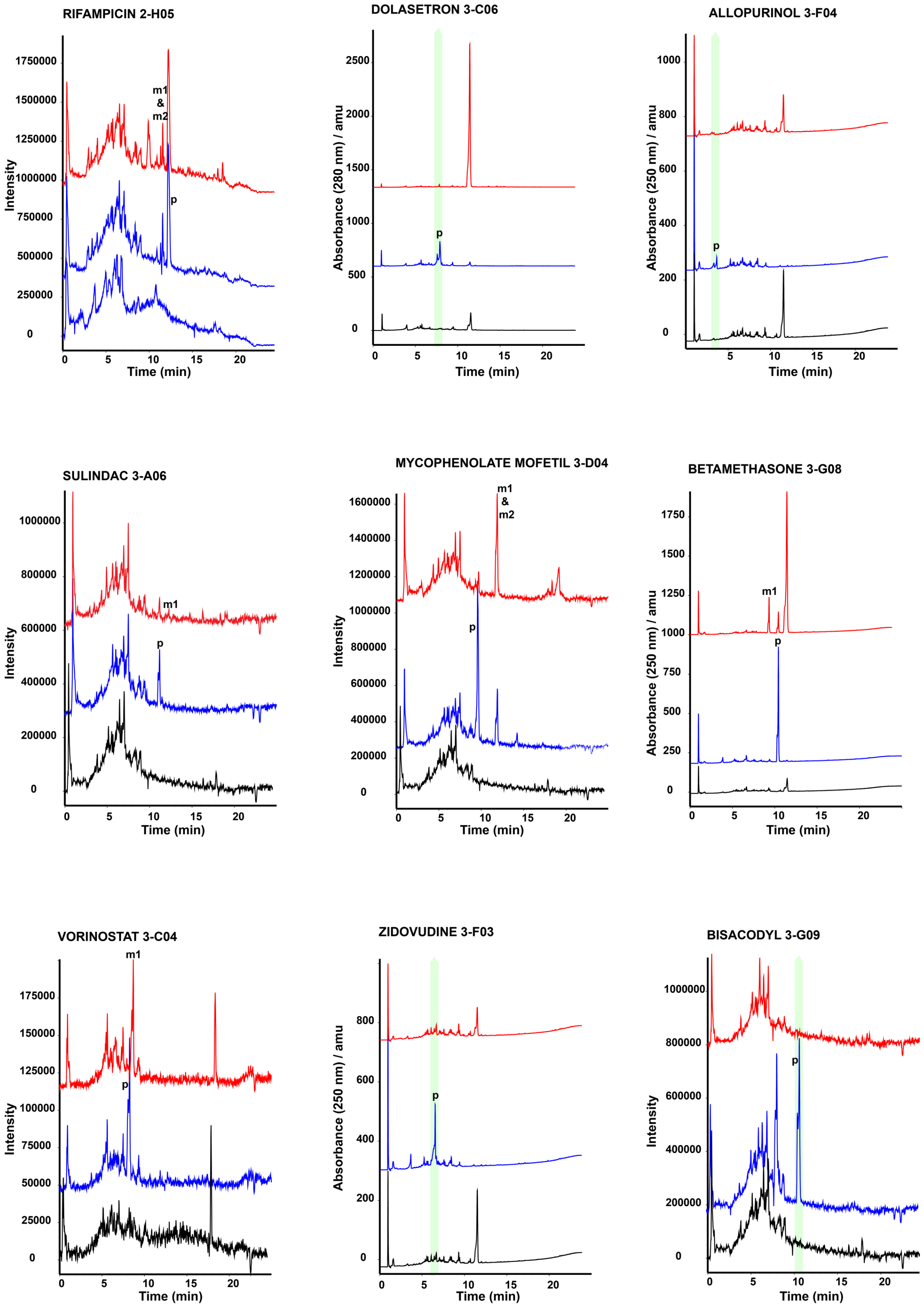
RIFAMPICIN 2-H05
DOLASETRON 3-C06
ALLOPURINOL 3-F04
SULINDAC 3-A06
MYCOPHENOLATE MOFETIL 3-D04
BETAMETHASONE 3-G08
VORINOSTAT 3-C04
ZIDOVUDINE 3-F03
BISACODYL 3-G09
Intensity
Absorbance (280 nm) / amu
Absorbance (250 nm) / amu
Time (min)
m1 & m2
m1
p

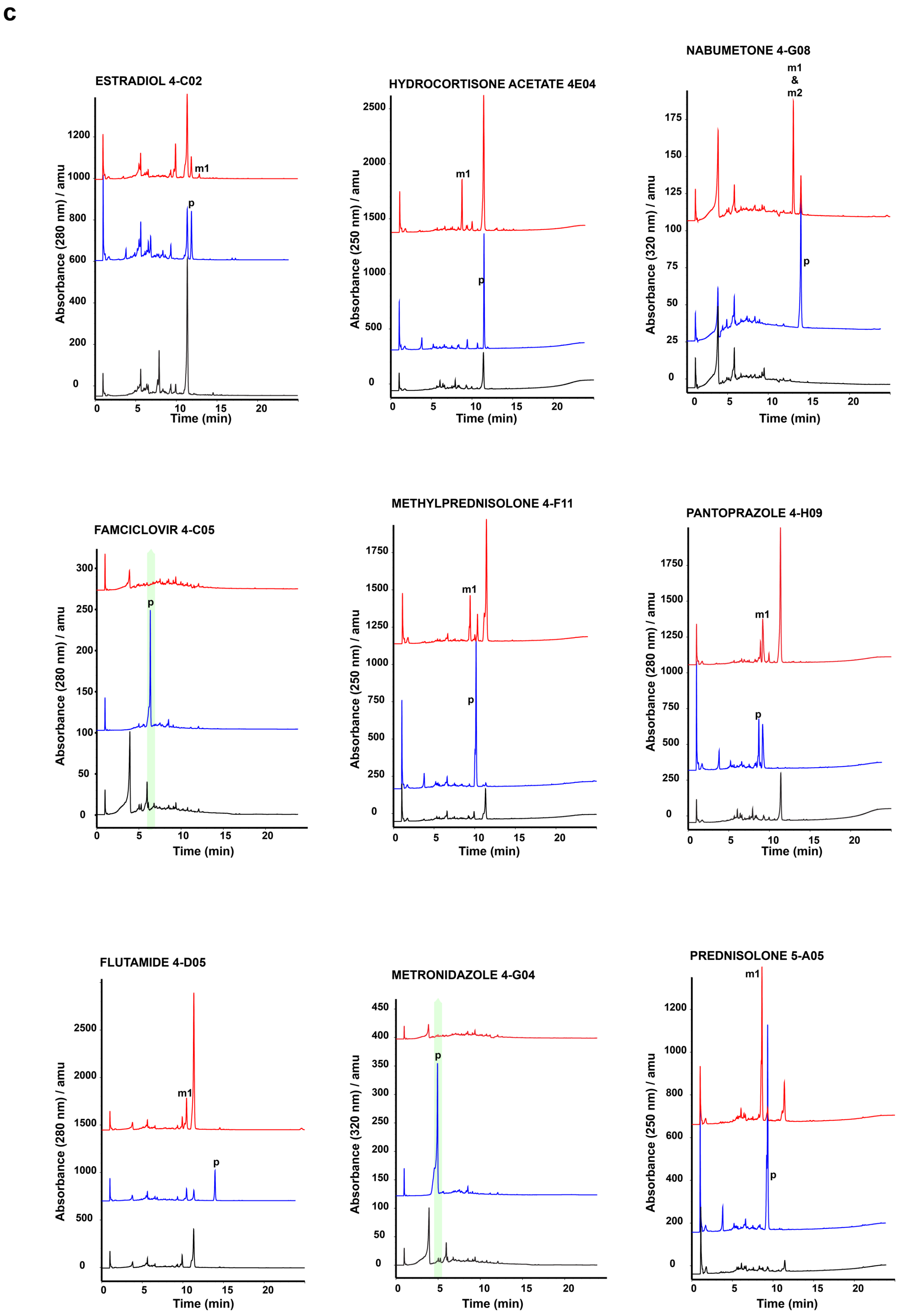
c
ESTRADIOL 4-C02
HYDROCORTISONE ACETATE 4E04
NABUMETONE 4-G08
FAMCICLOVIR 4-C05
METHYLPREDNISOLONE 4-F11
PANTOPRAZOLE 4-H09
FLUTAMIDE 4-D05
METRONIDAZOLE 4-G04
PREDNISOLONE 5-A05
Absorbance (280 nm) / amu
Absorbance (250 nm) / amu
Absorbance (320 nm) / amu
Time (min)
m1
m1 & m2
p

d

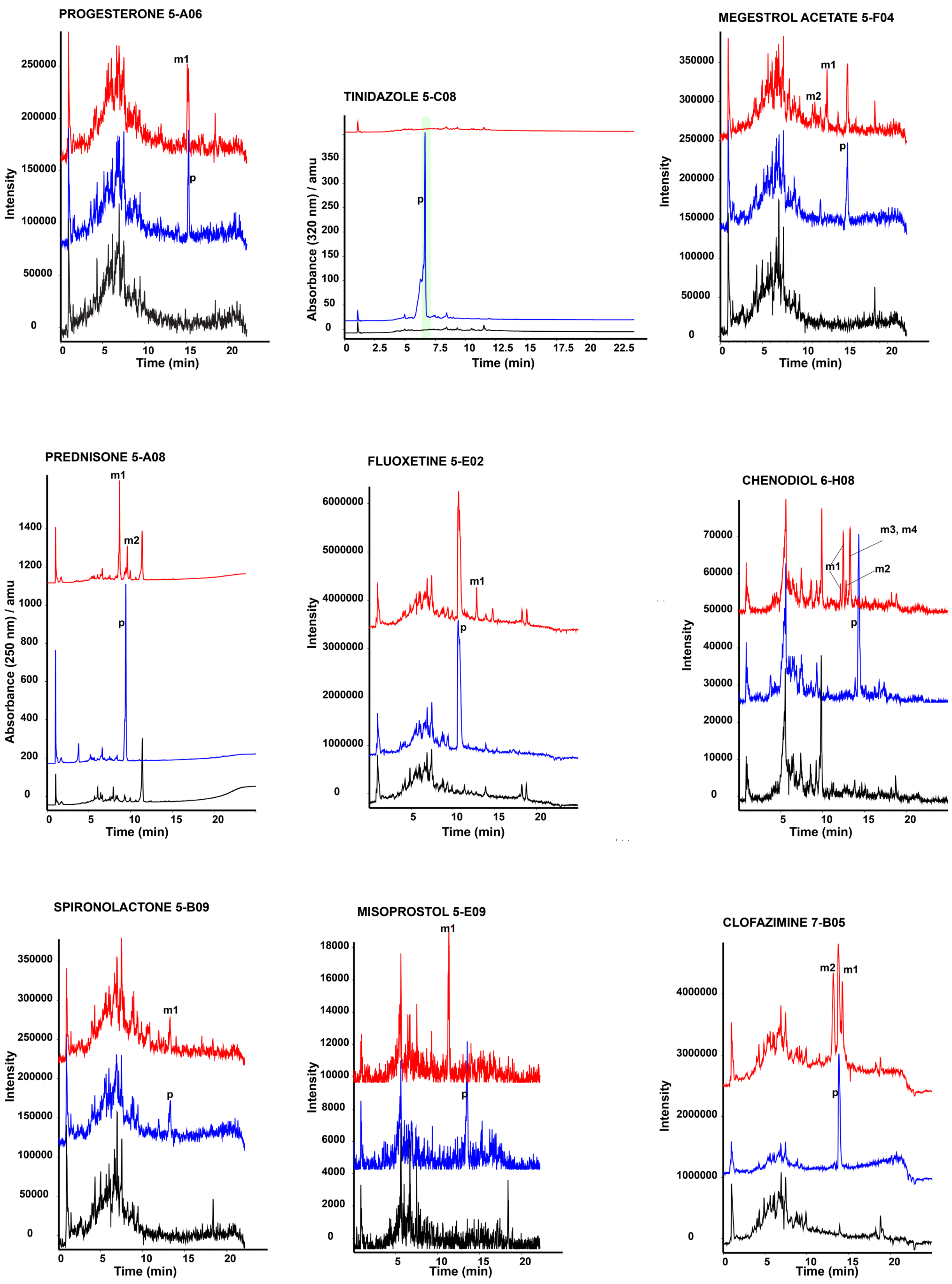
PROGESTERONE 5-A06
Intensity
Time (min)
m1
p
TINIDAZOLE 5-C08
Absorbance (320 nm) / amu
Time (min)
p
MEGESTROL ACETATE 5-F04
Intensity
Time (min)
m1
m2
p
PREDNISONE 5-A08
Absorbance (250 nm) / amu
Time (min)
m1
m2
p
FLUOXETINE 5-E02
Intensity
Time (min)
m1
p
CHENODIOL 6-H08
Intensity
Time (min)
m3, m4
m1
m2
p
SPIRONOLACTONE 5-B09
Intensity
Time (min)
m1
p
MISOPROSTOL 5-E09
Intensity
Time (min)
m1
p
CLOFAZIMINE 7-B05
Intensity
Time (min)
m2
m1
p

e

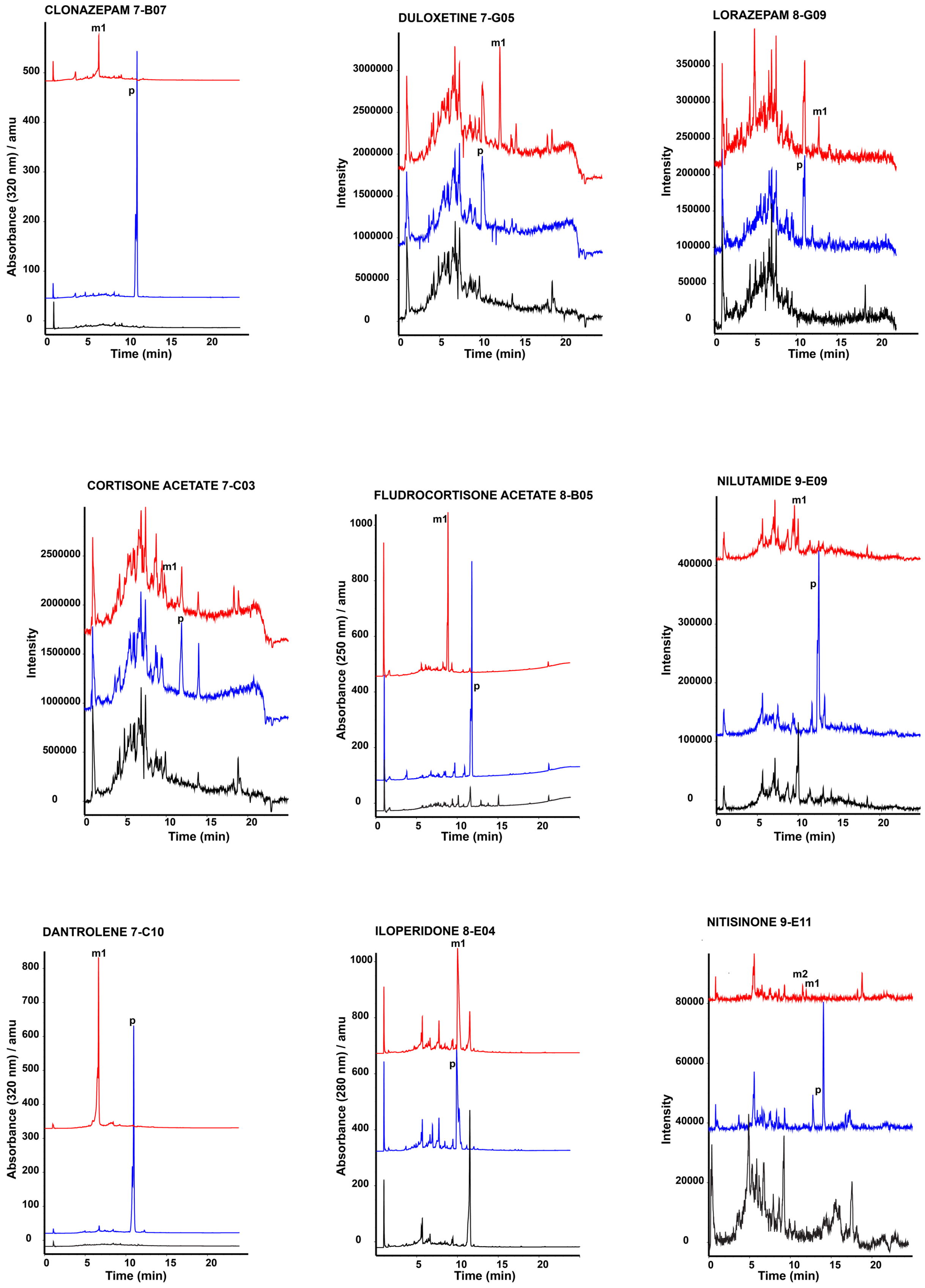

CLONAZEPAM 7-B07
m1
p
Absorbance (320 nm) / amu
0
100
200
300
400
500
0
5
10
15
20
Time (min)
DULOXETINE 7-G05
m1
p
Intensity
0
500000
1000000
1500000
2000000
2500000
3000000
Time (min)
LORAZEPAM 8-G09
m1
p
Intensity
0
50000
100000
150000
200000
250000
300000
350000
Time (min)
CORTISONE ACETATE 7-C03
m1
p
Intensity
0
500000
1000000
1500000
2000000
2500000
Time (min)
FLUDROCORTISONE ACETATE 8-B05
m1
p
Absorbance (250 nm) / amu
0
200
400
600
800
1000
Time (min)
NILUTAMIDE 9-E09
m1
p
Intensity
0
100000
200000
300000
400000
Time (min)
DANTROLENE 7-C10
m1
p
Absorbance (320 nm) / amu
0
100
200
300
400
500
600
700
800
Time (min)
ILOPERIDONE 8-E04
m1
p
Absorbance (280 nm) / amu
0
200
400
600
800
1000
Time (min)
NITISINONE 9-E11
m2
m1
p
Intensity
0
20000
40000
60000
80000
Time (min)

f

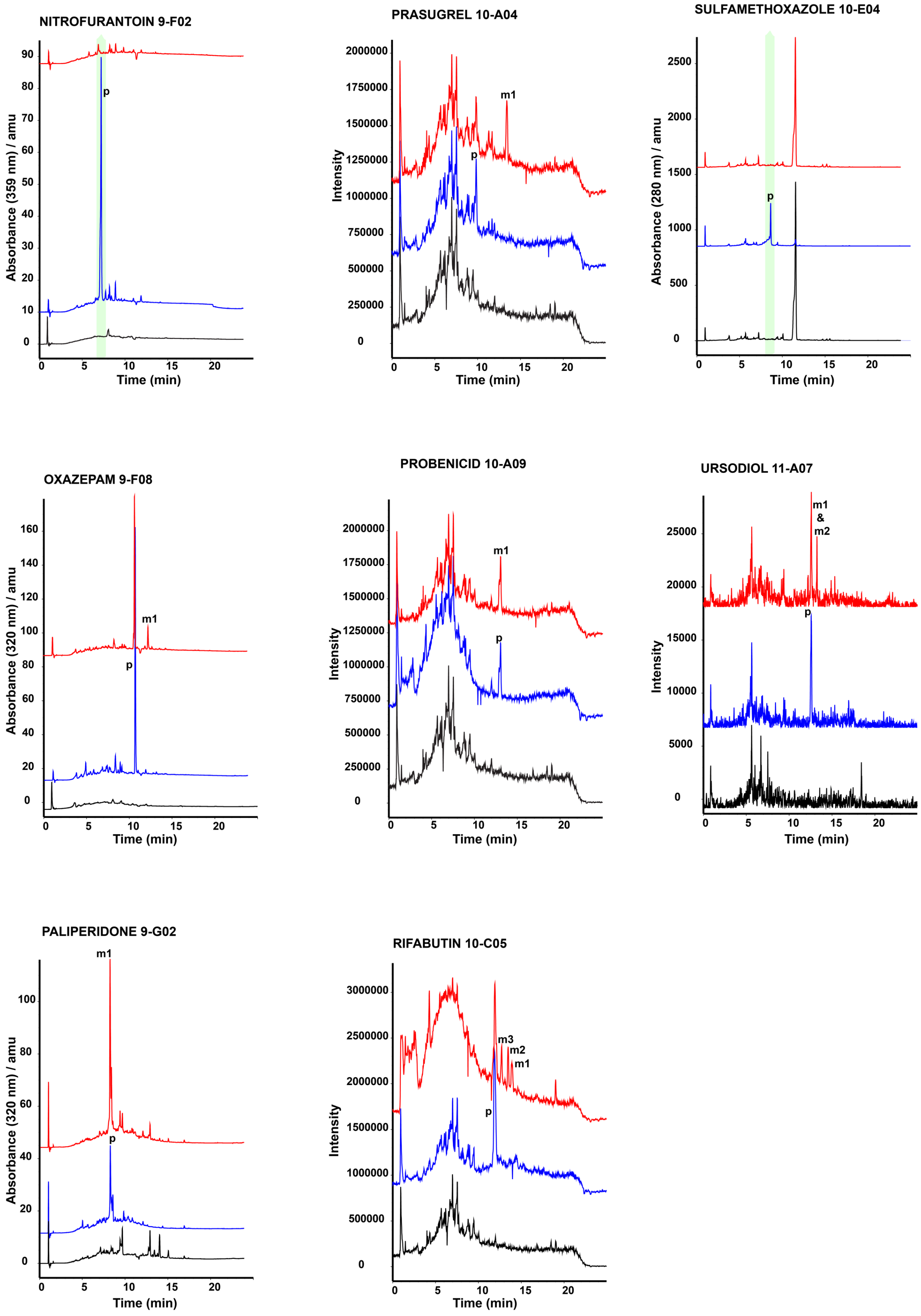
NITROFURANTOIN 9-F02
Absorbance (359 nm) / amu
Time (min)
p
PRASUGREL 10-A04
Intensity
Time (min)
m1
p
SULFAMETHOXAZOLE 10-E04
Absorbance (280 nm) / amu
Time (min)
p
OXAZEPAM 9-F08
Absorbance (320 nm) / amu
Time (min)
m1
p
PROBENICID 10-A09
Intensity
Time (min)
m1
p
URSODIOL 11-A07
Intensity
Time (min)
m1 & m2
p
PALIPERIDONE 9-G02
Absorbance (320 nm) / amu
Time (min)
m1
p
RIFABUTIN 10-C05
Intensity
Time (min)
m3
m2
m1
p

**Supplementary Figure 4. Sequential metabolism of hydrocortisone acetate by the HD-1 microbiome.** HPLC-MS analysis of hydrocortisone acetate (10) and hydrocortisone (6) incubated with HD-1 mGAM-02 culture, mGAM.02 broth, *P. distasonis* culture, or *C. bolteae* culture. An HPLC chromatogram at an absorbance of 254 nm is shown for all samples, indicating the conversion of hydrocortisone acetate to hydrocortisone by both *P. distasonis* and *C. bolteae*, and the conversion of both hydrocortisone acetate (in two steps) and hydrocortisone (in one step) to 20-dihydrocortisone (7) in the presence of the HD-1 microbiome.

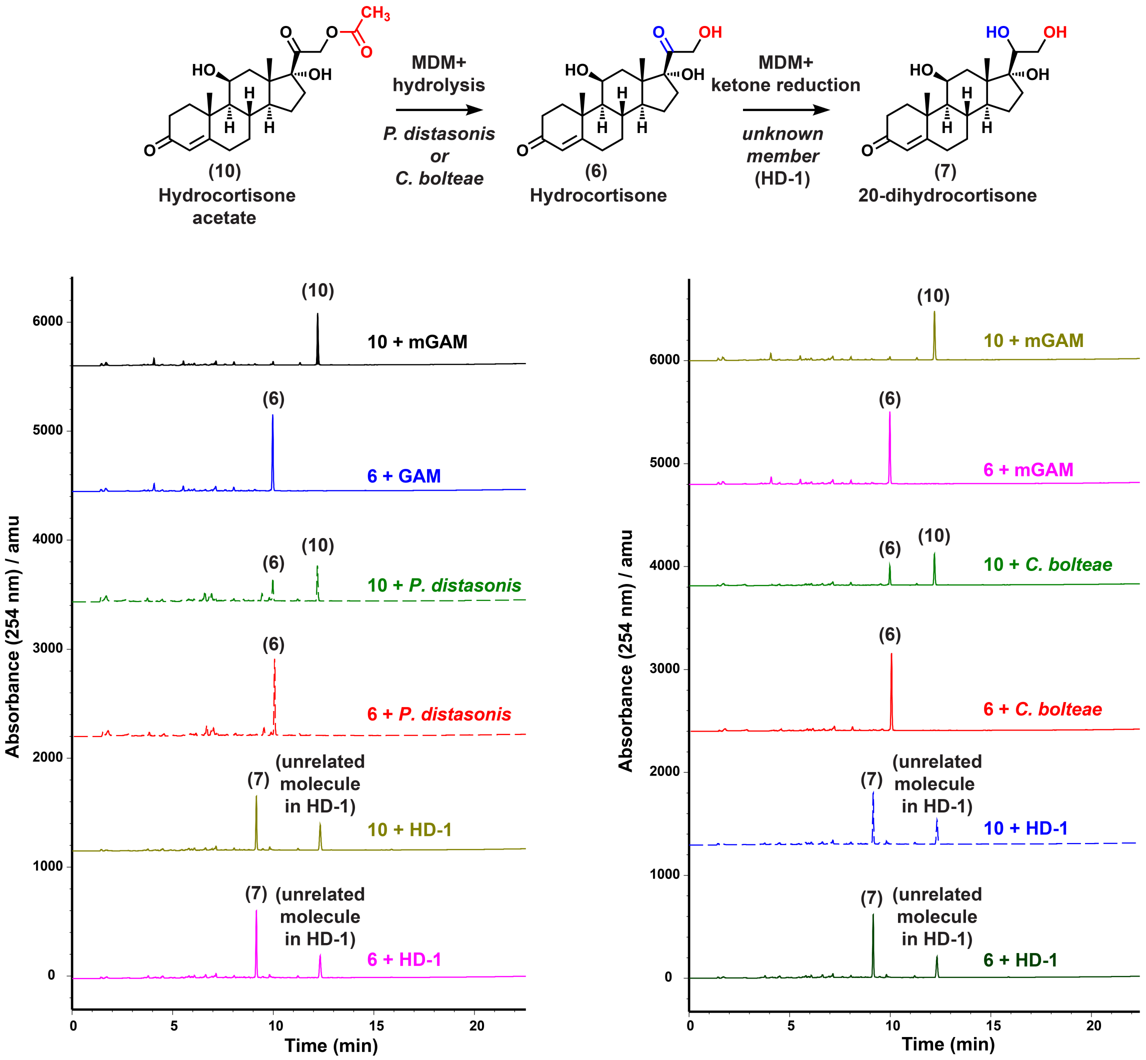

**Supplementary Figure 5. MDM deglycosylation of the anticancer drug trifluridine.** HPLC-MS analysis of trifluridine (17) incubated with wild type *E. coli* BW25113 (WT), and Δ*udp*, Δ*deoA*, and Δ*deoA*/Δ*udp* mutants in M9 medium. An HPLC chromatogram at an absorbance of 250 nm is shown for all samples, indicating the conversion of trifluridine (17) to trifluorothymine (18) by wild type *E. coli* BW25113 (WT), Δ*udp*, and Δ*deoA*, but not the Δ*deoA*/Δ*udp* mutant.

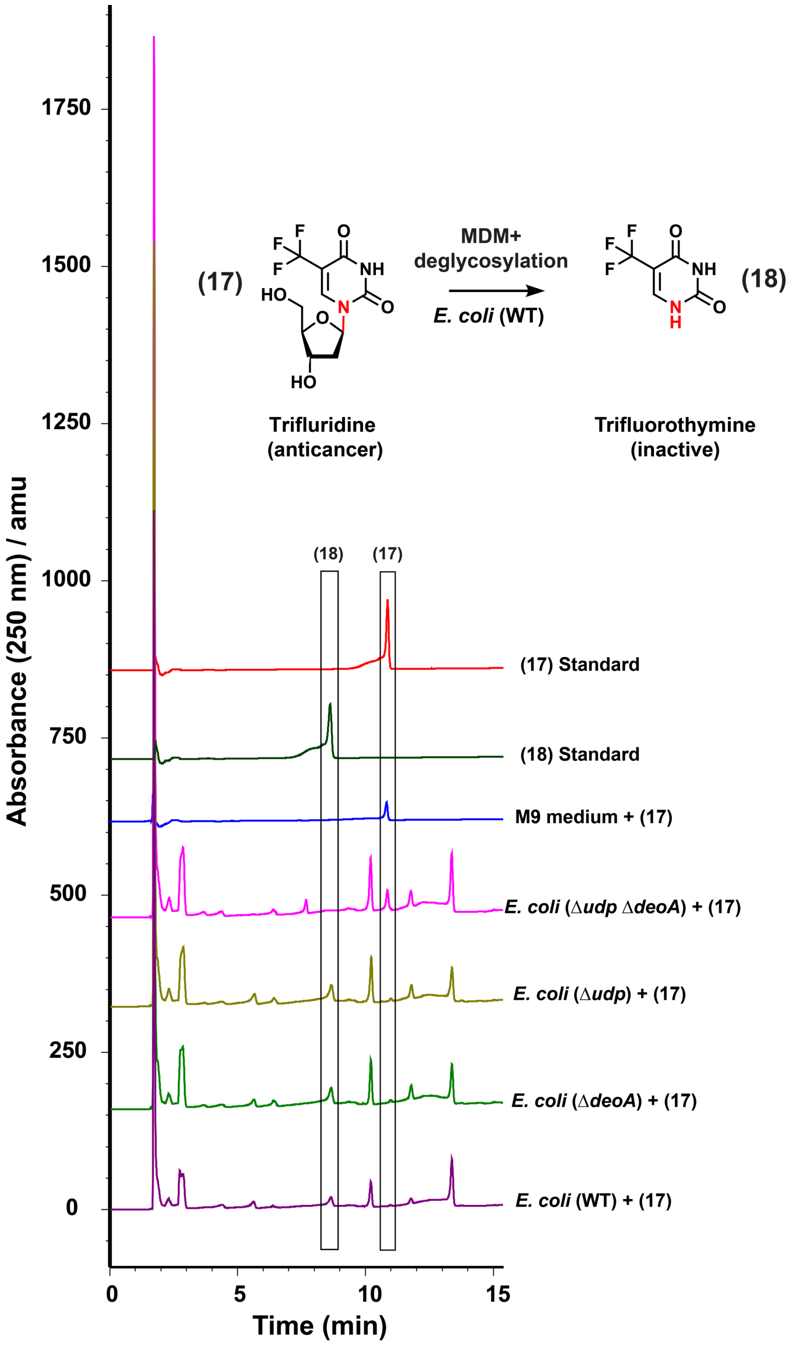

**Supplementary Figure 6. MDM deglycosylation of the anticancer prodrug doxifluridine.** HPLC-MS analysis of doxifluridine (19) incubated with wild type *E. coli* BW25113 (WT), and Δ*udp*, Δ*deoA*, and Δ*deoA*/Δ*udp* mutants in M9 medium. Extracted Ion Chromatograms for both doxifluridine (19) and its resulting MDM metabolite 5-fluorouracil (20) is shown for all samples, indicating the complete conversion of doxifluridine (19) to 5-fluorouracil (20) by wild type *E. coli* BW25113 (WT), Δ*udp*, and Δ*deoA*, but not the Δ*deoA*/Δ*udp* mutant.

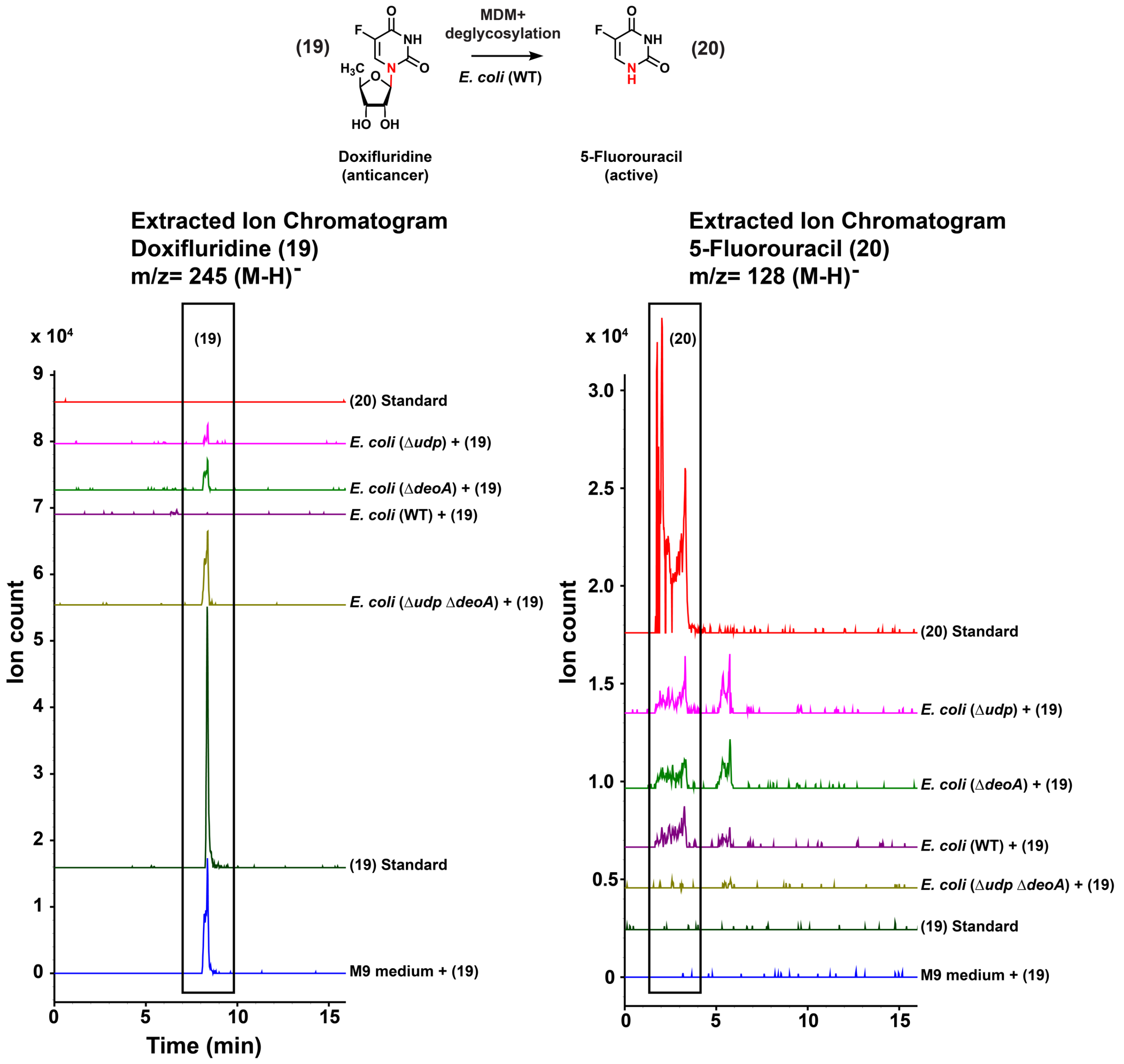

**Supplementary Figure 7. Capecitabine and deglycocapecitabine metabolism by human and bacterial enzymes. a)** HPLC-MS analysis of capecitabine (8) and deglycocapecitabine (9) incubated with human carboxyesterase 1 (CES), or heat inactivated human carboxyesterase 1 (HI CES). An HPLC chromatogram at an absorbance of 280 nm is shown for all samples, indicating the conversion of capecitabine (8) to 5'-Deoxy-5-Fluorocytidine (9) by CES. No change is observed to deglycocapecitabine (9) under the same conditions. **b)** Biochemical scheme for capecitabine metabolism by human enzymes reported previously (top) and the MDM arm discovered in this study (bottom).

**a**

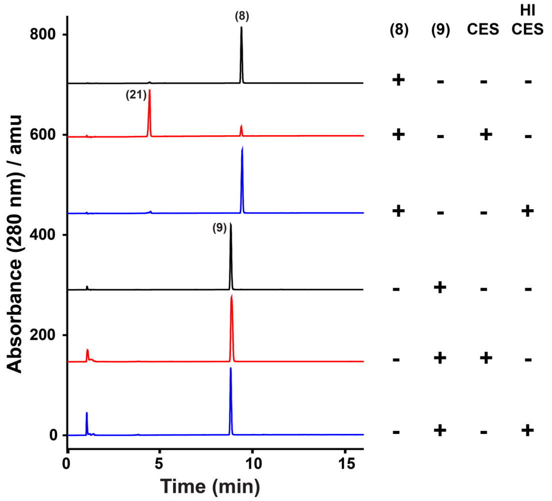


**b**

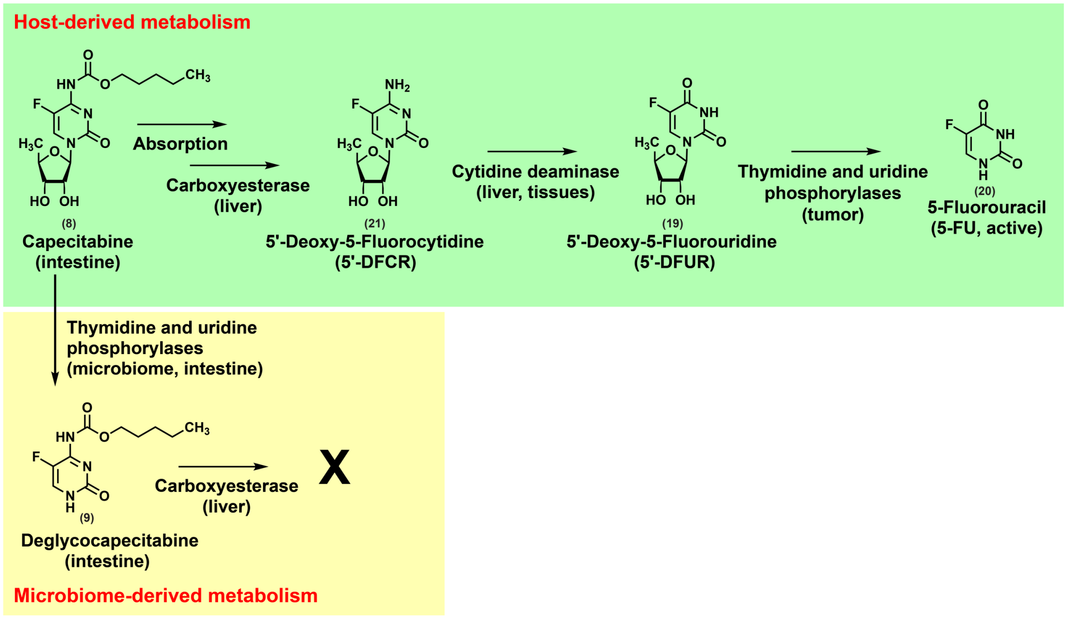

**Supplementary Figure 8. Microbiome-dependent pharmacokinetics of capecitabine.** HR-HPLC-MS based quantification of capecitabine and its human metabolite (5’-deoxy-5-fluorocytidine) in fecal and blood samples from mice colonized with HD-1 in comparison to non-colonized ones. No significant difference in the levels of capecitabine were observed between the two groups. ($p > 0.05$, significance was determined by testing the intersection null hypothesis with marginal two-tailed t-tests using the Bonferroni correction to control family-wise error rate). Error bars represent the standard error of the mean.

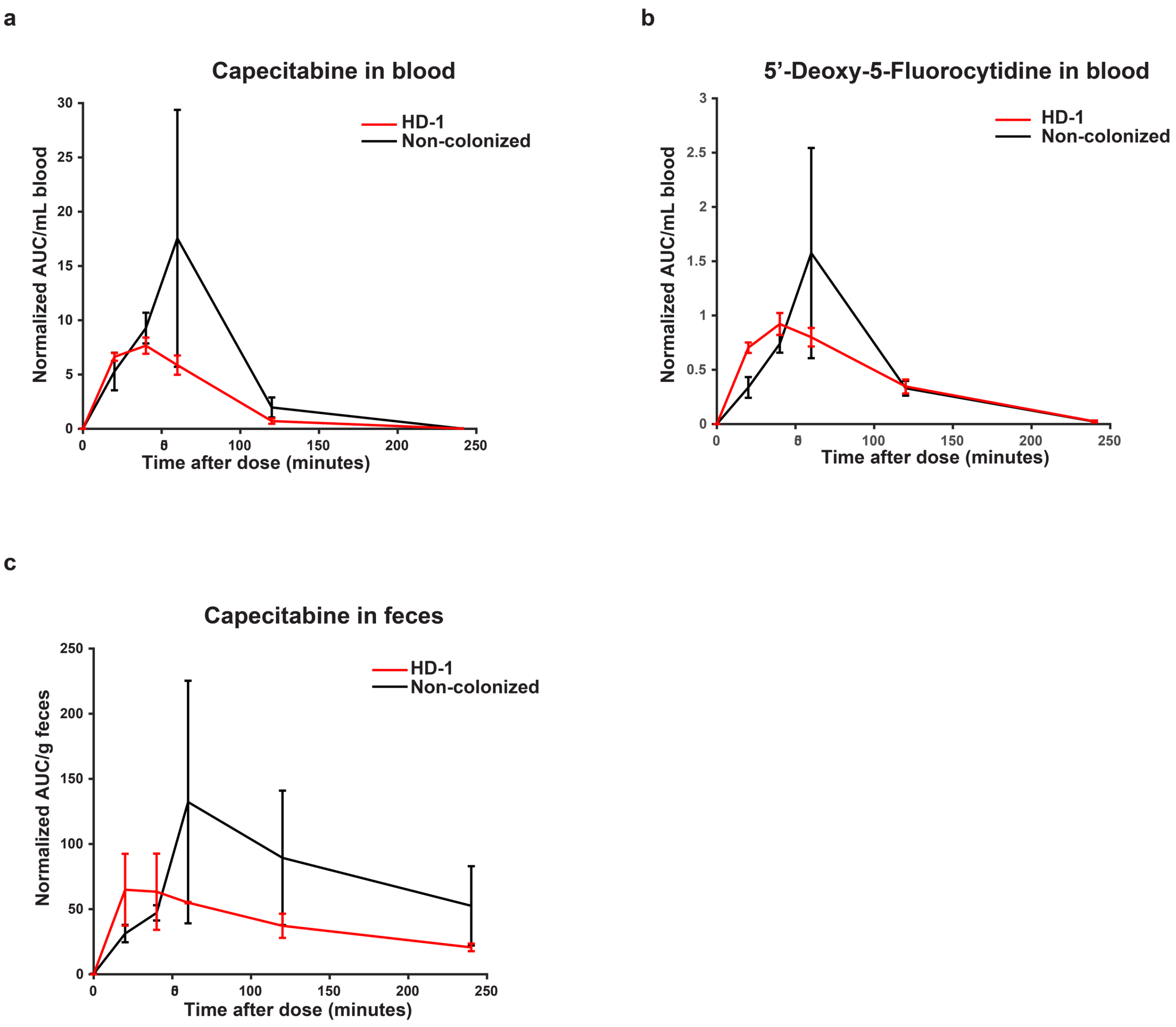
