## Supplementary Data 1 for "Systematic mapping of drug metabolism by the human gut microbiome"

**Structural determination of selected MDM metabolites**

**Table 1.** Molecular formulae of the purified metabolites determined by HRMS

| Compounds | predicted molecular formula | Calculate m/z $[M+H]^+$ | Observed m/z: $[M+H]^+$ |
|---|---|---|---|
| Aminonicardipine | $C_{26}H_{31}N_3O_4$ | 450.2393 | 450.2378 |
| 20-dihydrocortisone | $C_{21}H_{32}O_5$ | 365.2328 | 365.2311 |
| deglycocapecitabine | $C_{10}H_{14}FN_3O_3$ | 244.1097 | 244.1089 |

### 1.1 Capecitabine and deglycocapecitabine

The expected molecular formula of deglycocapecitabine ($C_{10}H_{14}FN_3O_3$) was deduced from HR-MS data (**Table 1**). To confirm its structure, we isolated pure deglycocapecitabine using HPLC. $^1$H-NMR spectrum of deglycocapecitabine was almost identical to that of capecitabine, except for the absence of the sugar moiety (**Table 2 and Figure 1**) and the presence of one proton of 1-NH at chemical shift 11.16 ppm. MS/MS experiments confirmed the absence of the sugar in deglycocapecitabine as compared to capecitabine (**Figure 2**). Altogether, the results confirmed the structure of the metabolite as deglycocapecitabine.

**Table 2.** $^1$H NMR data for capecitabine and deglycocapecitabine (500 MHz in DMSO-D6)

| Position | δH, mult (*J* in Hz) Capecitabine | δH, mult (*J* in Hz) Deglycocapecitabine |
|---|---|---|
| **1-NH** | ND* | 11.16, br |
| **2** | | |
| **3** | | |
| **4** | | |
| **5** | | |
| **6-CH** | 8.01, s | 7.95, s |
| **7-NH** | 10.52, br | 11.16, br |
| **8** | | |
| **9-$CH_2$** | 4.07, $m^b$ | 4.05, t |
| **10-$CH_2$** | 1.61, p | 1.60, p |
| **11-$CH_2$** | 1.32, $m^b$ | 1.32, $m^b$ |
| **12-$CH_2$** | 1.32, $m^b$ | 1.32, $m^b$ |
| **13-$CH_3$** | 0.88, t | 0.88, $m^b$ |
| **14-CH** | 5.67, d | ND |
| **15-CH**<br>**-OH** | 4.09, $m^b$<br>5.14 | ND |
| **16-CH**<br>**-OH** | 3.68, t<br>5.14, br | ND |
| **17-CH** | 3.90, p | ND |
| **18-$CH_3$** | 1.31, $m^b$ | ND |

*[b]* overlapped signal
*ND: Not Detected

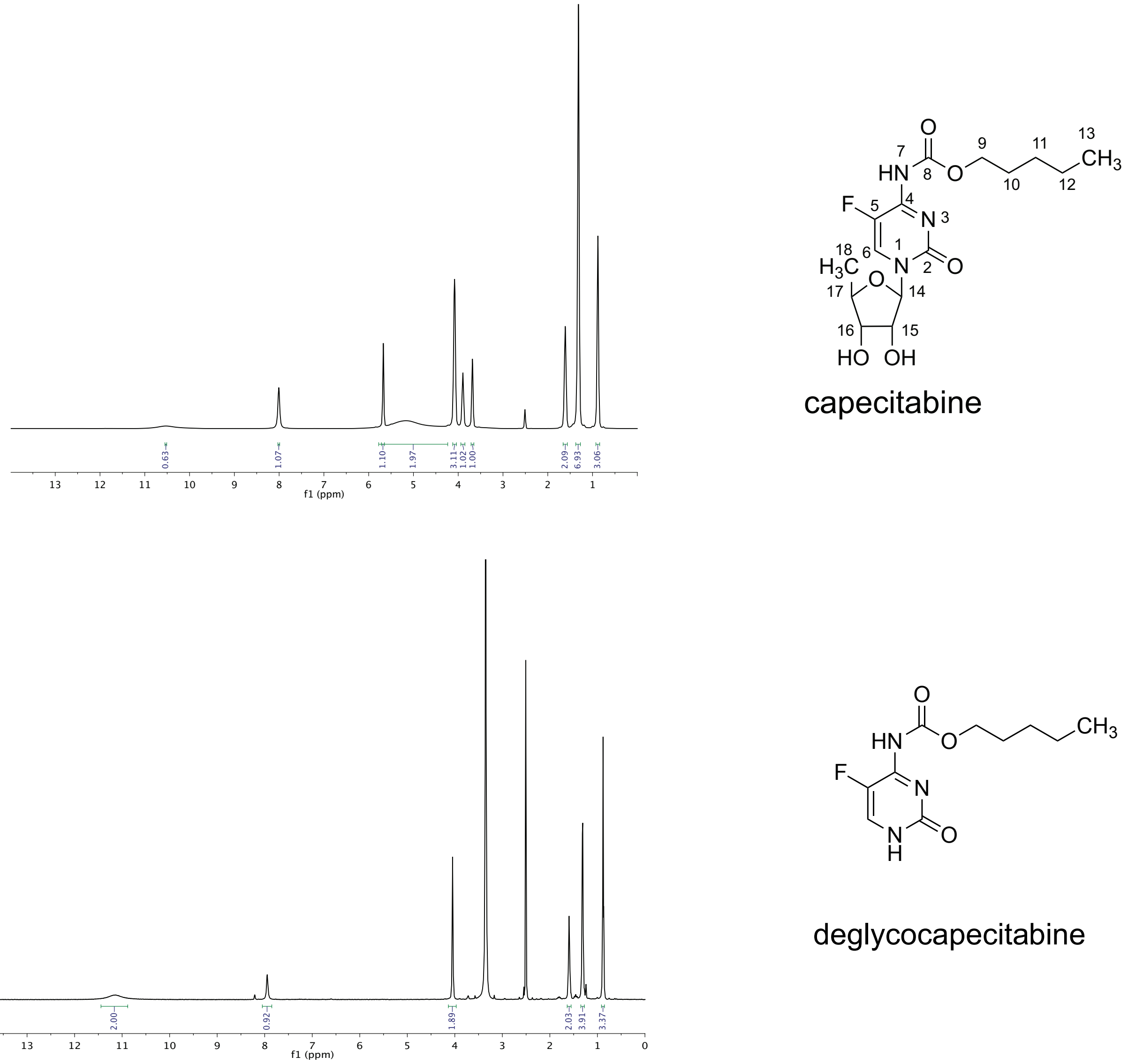


**Figure 1.** $^{1}$H NMR spectra of capecitabine and deglycocapecitabine in DMSO-D6

**HR-MS/MS of deglycocapecitabine** **HR-MS/MS of capecitabine**

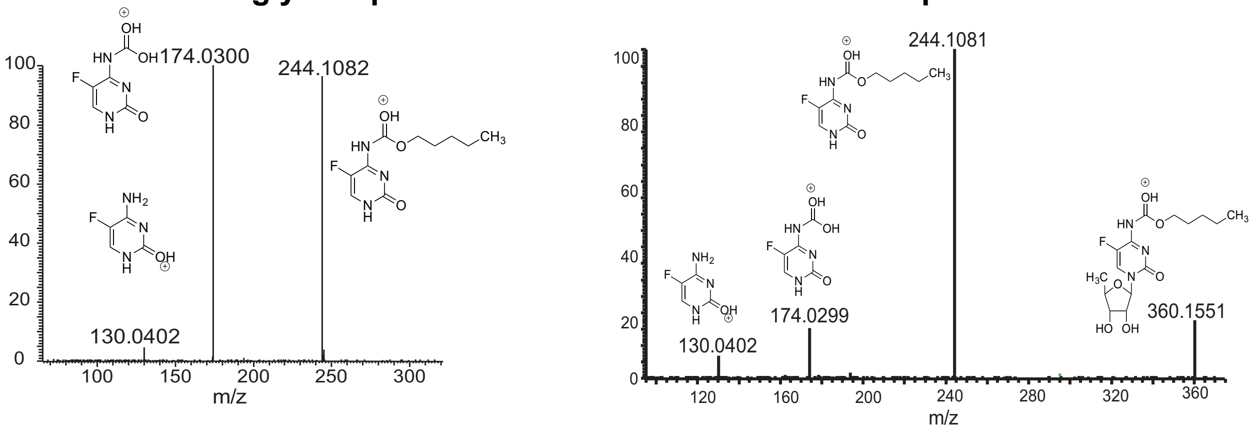


**Figure 2.** HR-MS/MS analysis of capecitabine and deglycocapecitabine

#### 1.2 Nicardipine and aminonicardipine

The molecular formula of aminonicardipine ($C_{26}H_{31}N_3O_4$) was deduced from HR-MS data (**Table 1**). To confirm its structure, we isolated pure aminonicardipine using HPLC. $^1$H-NMR spectrum of aminonicardipine was almost identical to that of nicardipine, except for the presence of two NH protons on C-9 at chemical shift of 8.51 ppm, and the upfield shift of the aromatic protons on C-8, 10, 11 and 12, which confirmed the absence of the electron withdrawing nitro group (**Table 3 and Figure 3**). Altogether, the results confirmed the structure of the metabolite as aminonicardipine, which derived from the nitroreduction of nicardipine.

**Table 3.** $^{1}$H NMR data for nicardipine and aminonicardipine (500 MHz in DMSO-D6)

| Position | δH, mult (*J* in Hz) Nicardipine | δH, mult (*J* in Hz) Aminonicardipine |
|---|---|---|
| **1-NH** | 9.40, s | 8.89 |
| **2** | | |
| **3** | | |
| **4-CH** | 5.00, s | 4.81, s |
| **5** | | |
| **6** | | |
| **7** | | |
| **8-CH** | 7.95, s | 7.24, m$^b$ |
| **9** | | |
| **9-NH$_2$** | ND* | 8.50, s$^b$ |
| **10-CH** | 7.95, s | 6.36, m$^b$ |
| **11-CH** | 7.60, m$^b$ | 6.79, m$^b$ |
| **12-CH** | 7.60, m$^b$ | 6.29, m$^b$ |
| **13-CH$_3$** | 2.34, s | 2.24, s |
| **14-CH$_3$** | 2.29, s | 2.13, s |
| **15** | | |
| **16-CH$_2$** | 4.42, m$^b$ | 4.10, s$^b$ |
| **17- CH$_2$** | 4.31,4.19, m$^b$ | 3.54, m$^b$ |
| **18-CH$_3$** | 2.56, s | 2.58, s$^b$ |
| **19-CH2** | 3.29, m$^b$ | 3.50, m$^b$ |
| **20** | | |
| **21-CH** | 7.45, m$^b$ | 7.29, m$^b$ |
| **22-CH** | 7.50, m$^b$ | 7.29, m$^b$ |
| **23-CH** | 7.45, m$^b$ | 7.29, m$^b$ |
| **24-CH** | 7.50, m$^b$ | 7.29, m$^b$ |
| **25-CH** | 7.45, m$^b$ | 7.29, m$^b$ |
| **26** | | |
| **27-CH$_3$** | 3.56, s | 3.50, m$^b$ |

$^b$ overlapped signal
*ND: Not Detected

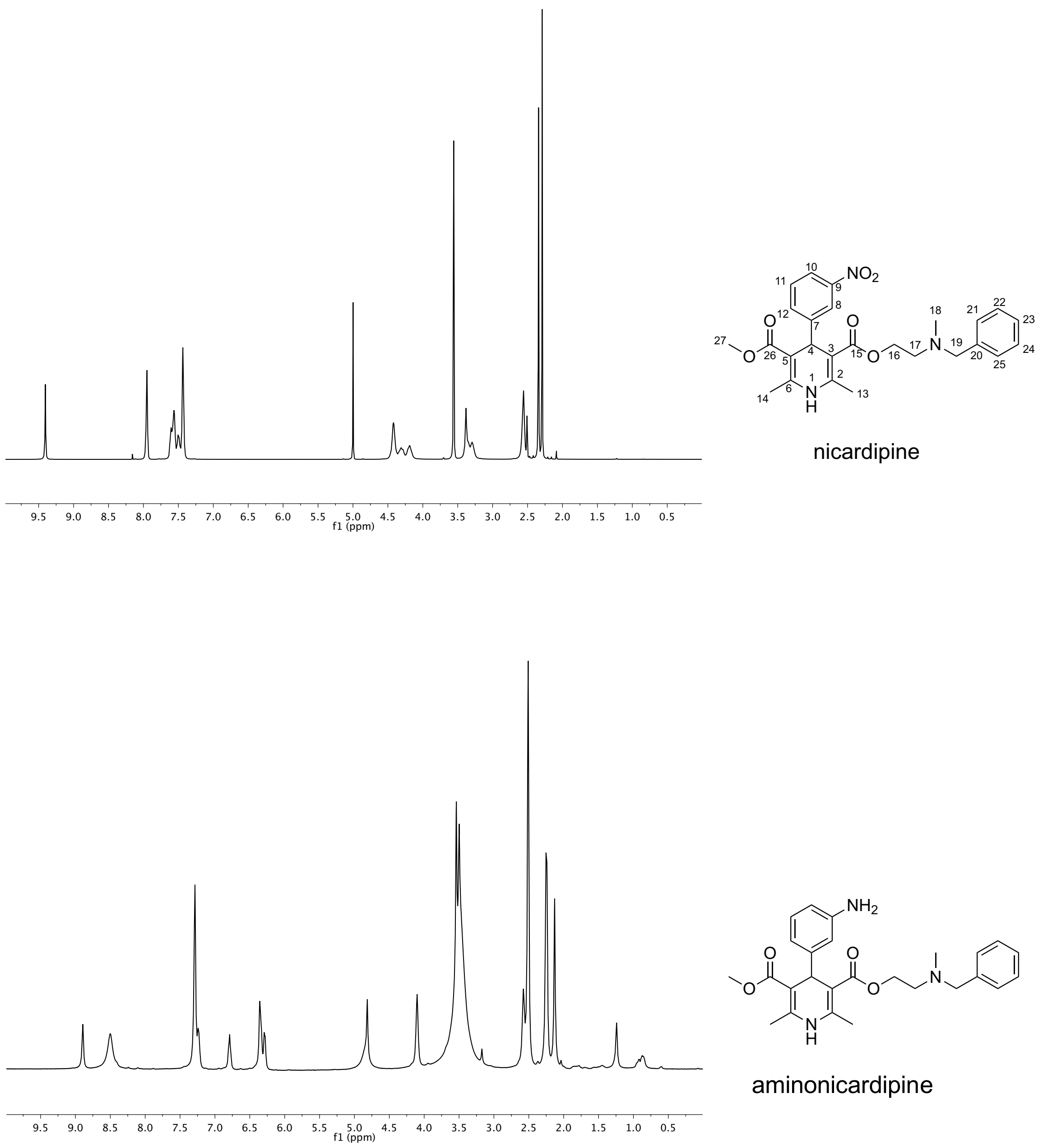


**Figure 3.** $^{1}H$ NMR spectra of nicardipine and aminonicardipine in DMSO-D6

#### 1.3 Hydrocortisone and 20-dihydrocortisone

The molecular formula of dihydrocortisone ($C_{21}H_{32}O_5$) was deduced from HR-MS data (**Table 1**), implying a reduction to the parent molecule. To elucidate the structure of the metabolite and localize the position of this modification, we isolated pure dihydrocortisone using HPLC. $^{13}C$ -NMR, $^{1}H$-NMR, and 2D-HSQC NMR spectra of dihydrocortisone were almost identical to those of hydrocortisone, except for: 1) The shift of the ketone carbon C-20 from chemical shift 211.7 ppm to 75.2 ppm, and the relocation of the methylene protons at C-21 from 4.08 and 4.51 ppm to 3.40 and 3.46 ppm, which resulted from the shielding effect of the hydroxyl group at C-20; and 2) The presence of a methine proton at C-20 at chemical shift 3.56 ppm (**Table 4 and Figures 4-6**). Altogether, the results confirmed the structure of the metabolite as 20-dihydrocortisone, which is derived from the ketone reduction of hydrocortisone.

**Table 4.** $^{1}H$ NMR data for hydrocortisone and 20-dihydrocortisone (500 MHz in DMSO-D6)

| Position | δH, mult (*J* in Hz) Hydrocortisone | δC | Position | δH, mult (*J* in Hz) 20-dihydrocortisone | δC |
|---|---|---|---|---|---|
| **1-$CH_2$** | 2.43, m$^b$ | 31.4 | **1-$CH_2$** | 1.32, 1.70 | 33.2 |
| **2-$CH_2$** | 2.57, t | 33.0 | **2-$CH_2$** | 2.43, 2.19 | 31.5 |
| **3** | | 198.1 | **3** | | 198.1 |
| **4-CH** | 5.56, s | 121.5 | **4-CH** | 5.55,s | 121.4 |
| **5** | | 172.4 | **5** | | 172.8 |
| **6-$CH_2$** | 2.41, 2.20, m$^b$ | 33.5 | **6-$CH_2$** | 2.18, 2.36 | 33.5 |
| **7-$CH_2$** | 1.65,1.27, m$^b$ | 23.4 | **7-$CH_2$** | | 23.8 |
| **8-CH** | 1.65, dd | 51.6 | **8-CH** | 1.5 | 50.3 |
| **9-CH** | 1.91, m$^b$ | 31.2 | **9-CH** | 1.87 | 31.3 |
| **10** | | 38.9 | **10** | | 38.9 |
| **11-CH** | 4.26, m | 66.5 | **11-CH** | 4.22, m | 66.7 |
| **12-$CH_2$** | 1.88, 1.54, m$^b$ | 39.1 | **12-$CH_2$** | 1.90, 1.63 | 40.9 |
| **13** | | 46.4 | **13** | | 46.5 |
| **14-CH** | 0.86, m | 55.6 | **14-CH** | 0.83, m | 55.8 |
| **15-$CH_2$** | 1.93, 0.99, m$^b$ | 32.8 | **15-$CH_2$** | 1.92, 0.95 | 33.0 |
| **16-$CH_2$** | 2.09,1.79, m$^b$ | 34.1 | **16-$CH_2$** | 2.10, 1.77 | 34.1 |
| **17** | | 88.5 | **17** | | 84.1 |
| **18-$CH_3$** | 0.75, s | 17.0 | **18-$CH_3$** | 0.98 | 17.1 |
| **19-$CH_3$** | 1.37, s | 20.5 | **19-$CH_3$** | 1.38 | 20.5 |
| **20** | | 211.7 | **20-CH** | 3.56, m$^b$ | 75.2 |
| **21-$CH_2$** | 4.51, 4.08, dd | 65.9 | **21-$CH_2$** | 3.45, 3.40, m$^b$ | 63.8 |

$^b$ overlapped signal

*ND: Not Detected

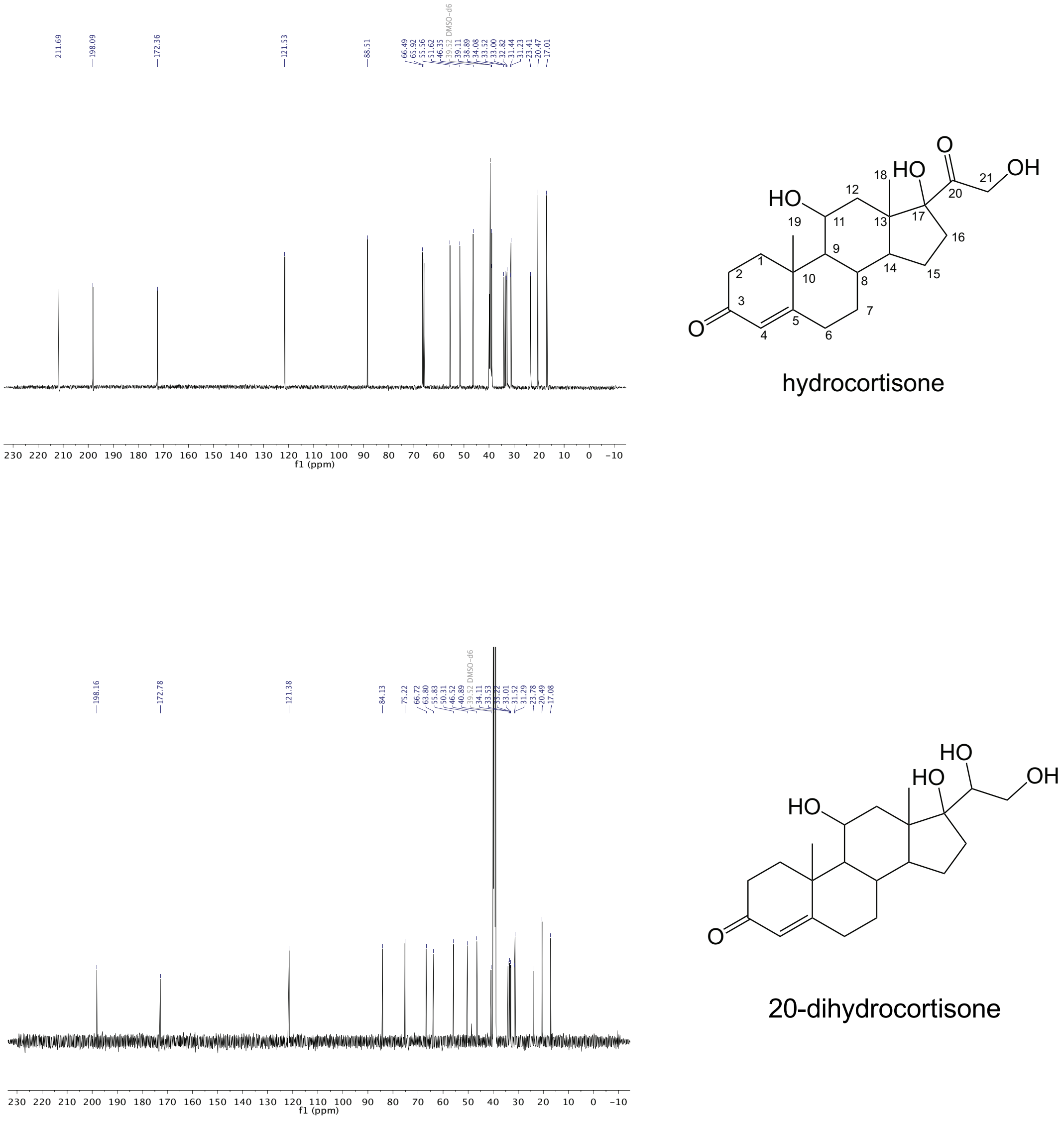


**Figure 4.** $^{13}$C NMR spectra of hydrocortisone and 20-dihydrocortisone in DMSO-D6

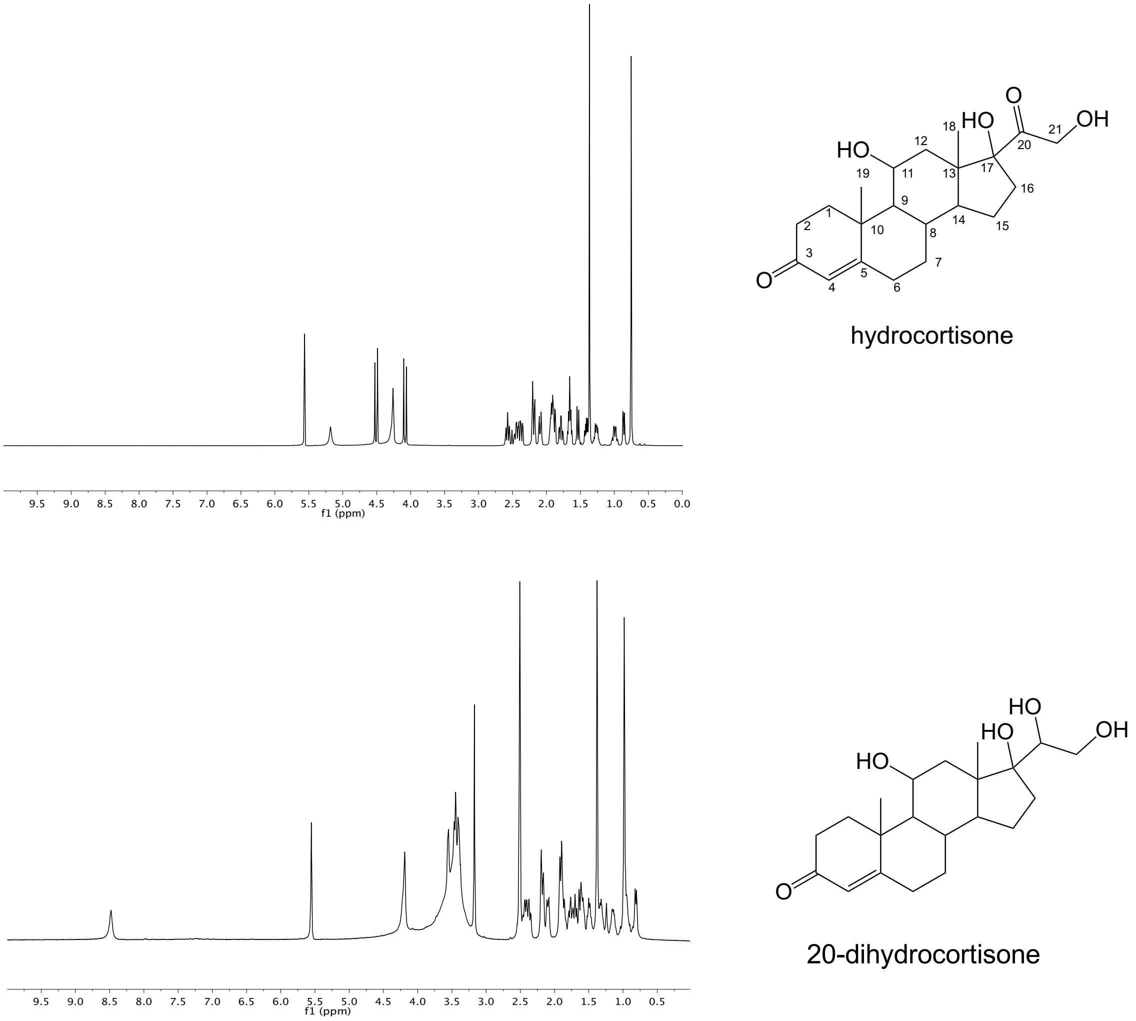


**Figure 5.** $^{1}H$ NMR spectra of hydrocortisone and 20-dihydrocortisone in DMSO-D6

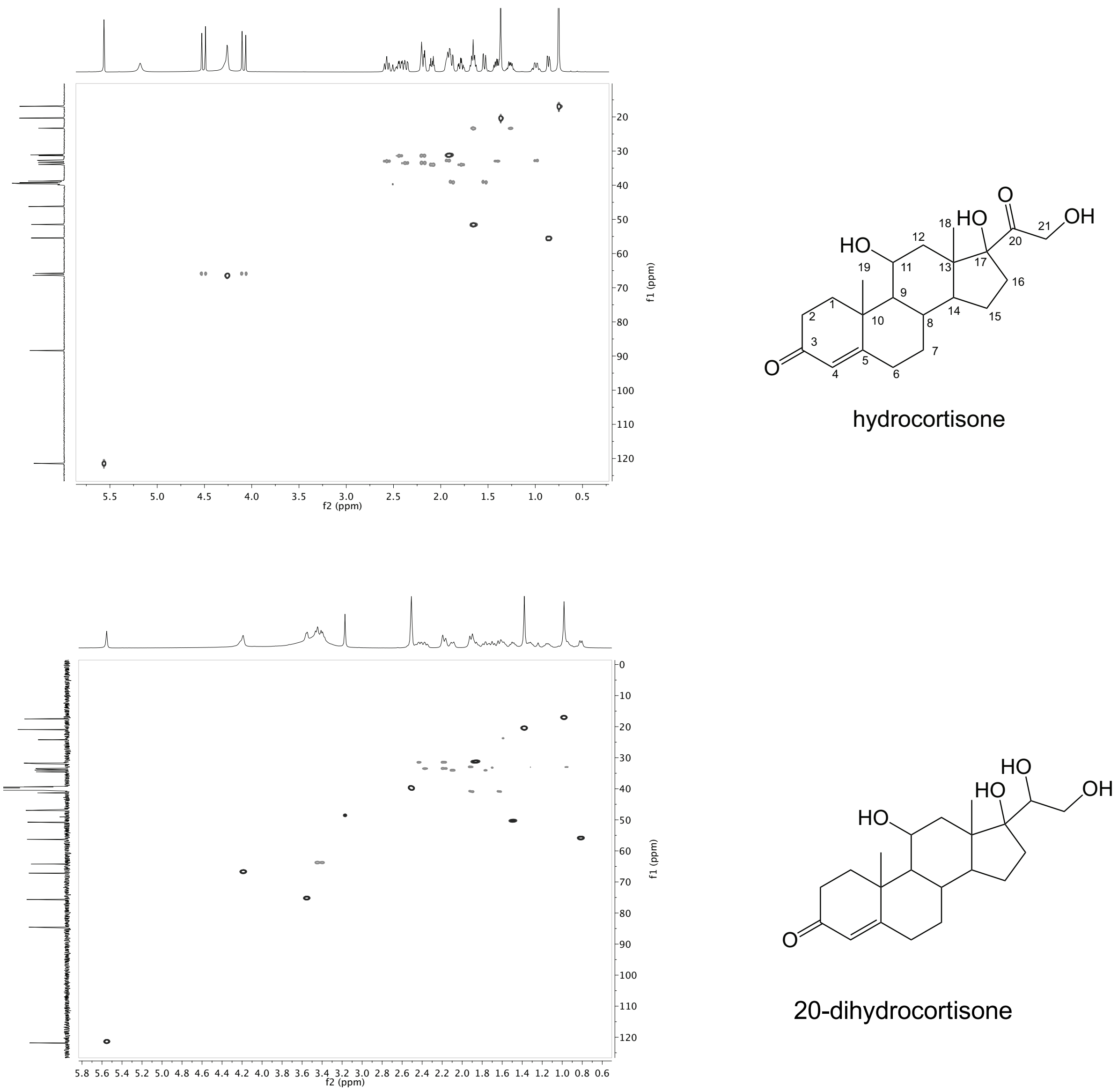


**Figure 6.** HSQC spectra of hydrocortisone and 20-dihydrocortisone in DMSO-D6
